# Neural abstraction of evidence accumulation enables flexible multidimensional perceptual decisions

**DOI:** 10.64898/2026.10.07.757231

**Authors:** Lucas Benjamin, Julie Drevet, Valentin Wyart

**Affiliations:** Laboratoire de Neurosciences Cognitives et Computationnelles (LNC^2^), Institut National de la Santé et de la Recherche Médicale (Inserm), Paris, France; Département d’Études Cognitives, École Normale Supérieure – Université PSL, Paris, France; Institut de Neurosciences des Systèmes (INS), Aix-Marseille Université, Marseille, France

## Abstract

Human perceptual decisions adapt to rich and changing contexts but are typically tested under impoverished conditions. To probe this flexibility, we examined decisions requiring both the selection and the combination of multiple sensory features across sequences of objects. Here we show that human decision flexibility extends beyond feature selection to combination, even across sensory modalities, supported by the abstraction of underlying neural representations. Rapid magnetoencephalographic (MEG) responses reveal a hierarchy of object representations that become increasingly abstract and gated according to their decision relevance. A shared abstract representation of updates reflects evidence accumulation collapsed onto the relevant axis across unidimensional (selection-based) and multidimensional (combination-based) contexts. Recurrent neural networks tested under the same conditions acquire and benefit from abstract representations only when operating under human levels of computational noise. Together, these findings provide a ‘flexibility through abstraction’ account of human decisions across a broader range of everyday perceptual conditions.

## Introduction

Human behavior is extremely flexible and context-sensitive. As an example, we routinely make accurate perceptual decisions from challenging and dynamically changing sensory surroundings. In everyday perception, this flexible behavior depends on multiple features of objects (e.g., their color, their shape, their motion, the sound they make, their texture) coming from multiple sensory modalities (in this case vision, audition, and touch). These multiple features must be selected, ignored or combined in a context-dependent manner, often on short timescales. However, research on perceptual decision-making has largely focused on unidimensional conditions that require processing a single object feature (Gold and Shadlen, 2007; Hanks and Summerfield, 2017; O’Connell and Kelly, 2021), or equates flexibility to the context-dependent selection of a unique relevant feature for subsequent accumulation (Okazawa and Kiani, 2023). While this **‘flexible selection’ account** captures an important aspect of decision-making, it does not explain how the brain combines two or more sensory features during multidimensional – including multimodal – perceptual decisions, arguably the norm rather than the exception in everyday perception. An alternative account of decision flexibility proposes that evidence is not accumulated in the stimulus (input) space defined by decision-relevant sensory features, but instead in the action (output) space defined by the effectors (e.g., saccades, hand movements, navigation) (Cisek and Kalaska, 2010; Shadlen and Kiani, 2013; Shushruth et al., 2018). This **‘flexible sensorimotor mapping’ account** effectively decouples evidence accumulation from the sensory features on which it is based, but ties it to action planning (Donner et al.,2009; van den Brink et al., 2023), precluding covert perceptual decisions that are not readily translated onto behavior.

Here we test and validate a third account of decision flexibility, in which evidence is accumulated in an intermediate, abstract representational space that is decoupled both from upstream sensory representations and downstream effector representations. This **‘flexibility through abstraction’ account** posits that this abstract space support not only the selection and accumulation of evidence about a single relevant sensory feature, as shown by previous research (Bennur and Gold, 2011; O’Connell et al., 2012; Mante et al., 2013; Pagan et al., 2025), but also the combination of sensory features during multidimensional perceptual decisions. Below, we first probe human decision flexibility across a wide range of contexts by asking participants to categorize sequences of colored shapes according to a frequently changing context (**Exp. 1**: visuo-visual decisions; **Figure 1A**; see **Methods**). Depending on the active context, categories were defined either by a single object feature (e.g., the ‘red’ category in the 1D color context) or by combinations of the two object features (e.g., the ‘blue square’ category in the 2D context), and were associated with overlapping probability distributions in the two-dimensional object space (**Figure 1B**). Sequences were engineered by sampling objects from category-specific probability distributions, such that accurate decisions required accumulating evidence from successive objects about about the category of the sequence according to the active context. Importantly, the mapping between categories and effectors (left- vs right-hand movements) could change from one trial to the next and was revealed only at response time, preventing evidence accumulation in action space. We further examine multimodal perceptual decisions using a variant of the same paradigm using sequences of audio-visual objects comprising a visual feature (color) and an auditory feature (pitch) (**Exp. 2**: audio-visual decisions; **Figure 1G**).

**Figure 1.**
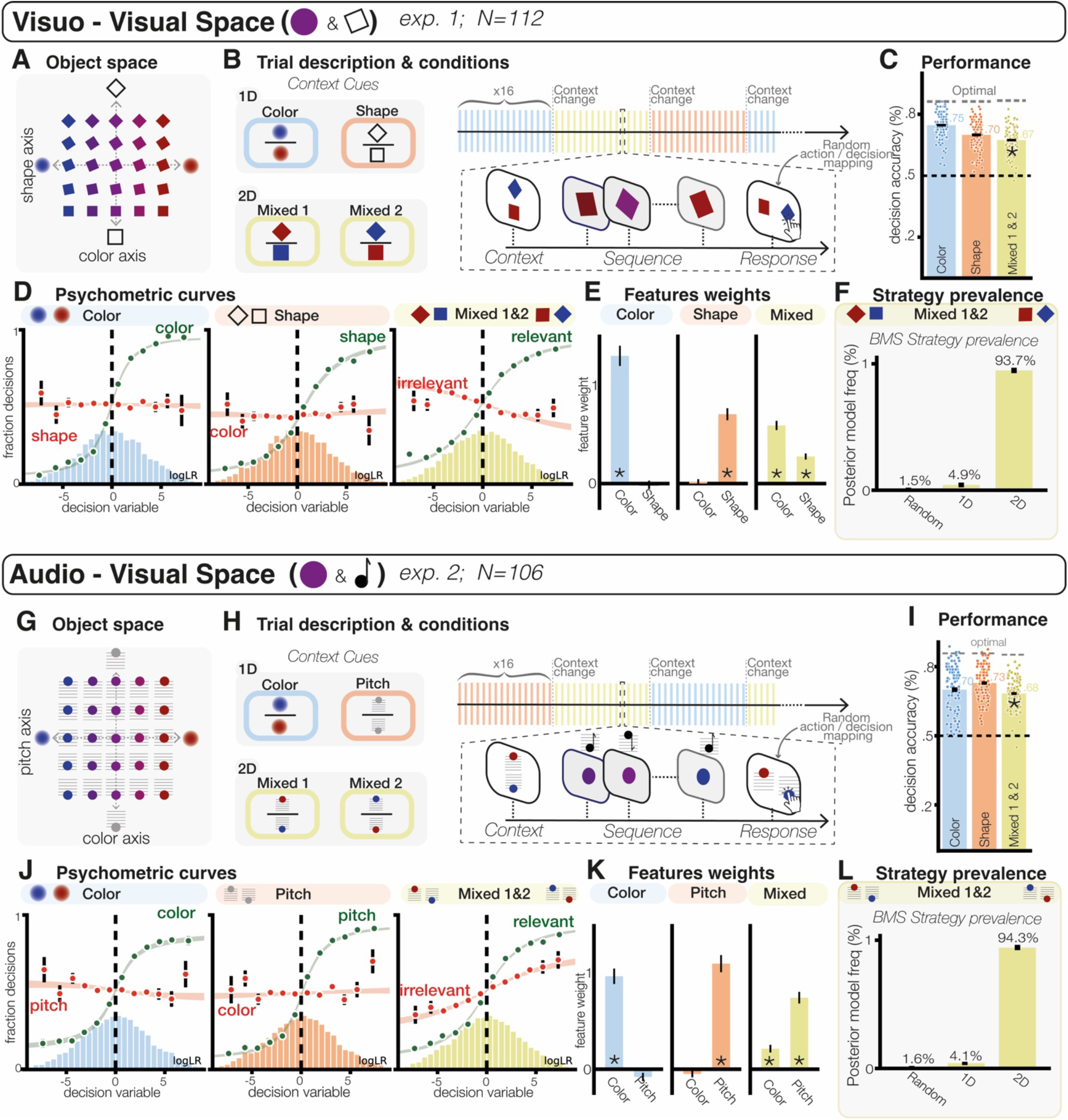
Task design and behavioral performance. **(A&G)** Objects spanned a two-dimensional space varying in color (from blue to red) and in shape (from square to diamond) for Exp. 1 or in pitch (low to high) in Exp. 2. **(B&H)** Left: context cues (top: 1D contexts; bottom: 2D contexts). Right: block and trial description. The active context changed every 16 trials. **(C&I)** Human categorization accuracy for each condition: color, shape (pitch), and mixed (2D) contexts. **(D&J)** Behavioral metrics. Left: color context. Center: shape/pitch context. Right: mixed (2D) contexts. Psychometric curves showing the relation between the decision variable (the accumulated category evidence in the active context) for the relevant (green) and irrelevant (red) feature, and the fraction of decisions. Dots and error bars indicate human data (mean ± s.e.m.) and shaded areas indicate psychometric fits (mean ± s.e.m.). **(E&K)** Left: color context. Center: shape/pitch context. Right: mixed (2D) contexts. Weight of each feature for the decision of the subjects in each condition. **(F&L)** Comparision of strategy prevalence in 2D conditions between a random strategy, a 1D strategy selecting a single object feature on individual trials, and the 2D strategy combining the two object features on individual trials. One star indicates a significant effect at *p* < 0.05.

Our empirical demonstration of **‘flexibility through abstraction’** proceeds in two steps. Behavioral results (**Exp. 1-2**) first establish that participants flexibly combine object features on every trial, rather than alternating between individual features across trials. Rapid MEG responses (**Exp. 3**) then reveal how this decision flexibility is implemented: the human brain selects and combines sensory features by projecting successive objects onto an abstract category axis before accumulating evidence, rather than accumulating evidence for each feature separately. This flexible abstraction allows for the reuse of the same neural representation to accumulate evidence across 1D and 2D contexts.

## Results

**Human behavior** To assess the accuracy of multidimensional visual decisions, we first compared human categorization accuracy between 1D (color and shape) and 2D (mixed) contexts (**Exp. 1**; **Figure 1** and **Figure S1**), and found that participants could perform 2D decisions nearly as well as equally difficult 1D decisions (group-level means, color: 75%, shape: 70%, mixed: 67%; **Figure 1C**). In 1D contexts, participants decided solely based on the relevant feature of sampled objects (e.g., pitting ‘red’ against ‘blue’ in the color context) and ignored the irrelevant feature (**Figure 1DE**), indicating efficient feature selection. In 2D contexts, the same participants made decisions based on the relevant combination of object features (e.g., pitting ‘blue square’ against ‘red diamond’), with close-to-zero interference from the irrelevant combination (**Figure 1DE**). Moreover, participants could adapt their decisions to the frequent changes in context with zero lag, reaching asymptotic accuracy from the first sequence after each context change. Critically, decisions made in 2D contexts reflected the combination of object features rather than the average of 1D decisions on individual trials. Indeed, Bayesian model selection (BMS) favored a strategy combining color and shape over a competing strategy alternating between 1D decisions about color or shape (estimated prevalence: 93.7%, exceedance *p* > 0.999; **Figure 1F**).

To test whether this combination strategy extends beyond object features from the same sensory modality, we created a variant of our multidimensional categorization task using audio-visual objects – beeping color flashes (**Exp. 2**; **Figure 1GH**). Participants could perform audio-visual 2D decisions about audio-visual objects nearly as well as equally difficult 1D decisions (group-level means, color: 70%, auditory pitch: 73%, mixed: 68%; **Figure 1IJK**). Similarly to what was found for visual 2D decisions, audio-visual 2D decisions decisions reflected the combination of color and auditory pitch rather than the average of 1D decisions about color or auditory pitch on individual trials (estimated prevalence: 94.3%, exceedance *p* > 0.999; **Figure 1L**).

### Neural selection of object features

To understand how the brain selects and combines multiple object features, we turned **Exp. 1** (visuo-visual decisions) into a magnetoencephalography (MEG) experiment (**Exp. 3**). To measure selection of the context-relevant feature in 1D contexts (color and shape), we analyzed MEG responses to each object of the sequence, by decoding three properties associated with object processing: 1. the **aspect** (color and shape) of the object, 2. the amount of color and shape **evidence** provided by the object, and 3. the **update** (increase or decrease) in accumulated evidence triggered by the object (Weiss et al., 2021) (**Figure 2A**; see **Methods**). We decoded these three properties from spatial patterns of MEG activity aligned to object presentation onset, separately for the context-relevant feature (color or shape) and the context-irrelevant feature. For the context-relevant feature, this analysis revealed a temporal sequence of decodability (Siegel et al., 2015; Brincat et al., 2018; Gwilliams and King, 2020; King and Wyart, 2021) for these three object properties (cluster-level decodability: all *p* < 0.001; **Figure 2B**, top), from the aspect of the object peaking at 180 ms following object presentation, the evidence provided by the object at 200 ms, and the update triggered by the object at 310 ms (peak latency differences: all *p* < 0.05; **Figure S3**). Decodability spread spatially along the dorsal pathway, from early visual cortex for the color and shape of the object to widespread associative regions including the posterior parietal cortex (PPC) and the dorsolateral prefrontal cortex (DLPFC) for the update triggered by the object (**Figure 2B**, bottom). Each object property was associated with highly dynamic neural representations – i.e., changing spatial patterns of MEG activity across time (**Figure S4**). We measured the strength of **selection** at each object processing stage (aspect, evidence, update) by computing the difference in decodability between context-relevant and context-irrelevant features. We found significant feature selection at all object processing stages (cluster-level difference: all *p* < 0.001; **Figure 2A**). Importantly, the strength of feature selection increased across processing stages, from a moderate 23% selection at the aspect stage to near-perfect 96% selection at the update stage. At this last stage, only the context-relevant update could be decoded better than chance from MEG activity.

**Figure 2.**
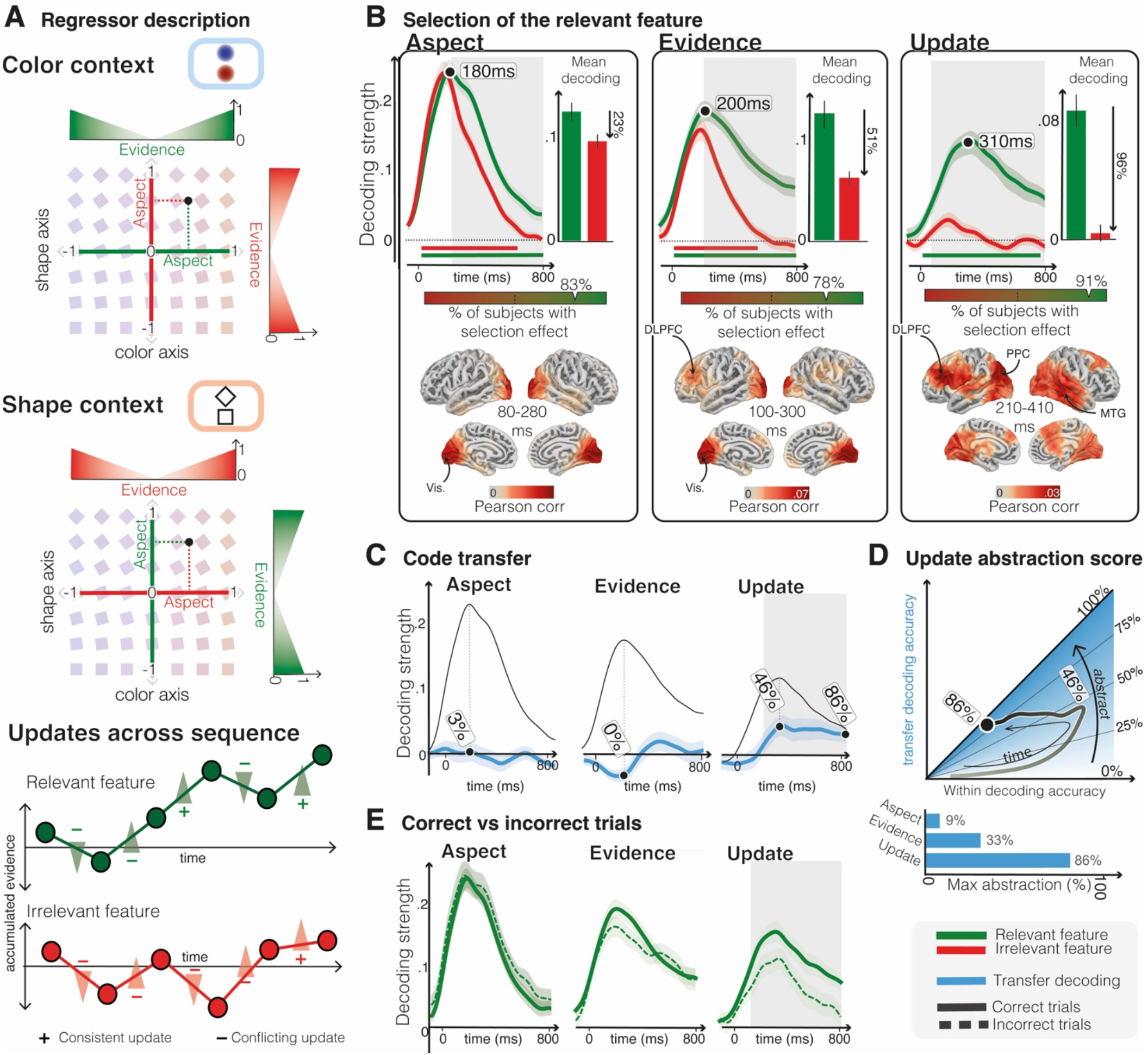
Neural selection and abstraction of object features in 1D contexts. **(A)** Description of the different object properties studied (see methods), **(B)** Decoding of the three different properties (aspect, evidence & update) for the relevant (green) and irrelevant (red) features. Significant clusters (cluster-based permutation analysis) are indicated with lines at the bottom of the graph (p<0.05). Significance between relevant and irrelevant features decoders (cluster-based permutation analysis) are shaded in grey (all p<0.001). Vertical bars represent the average decoding value over the significant window. Bottom: source localization and searchlight ridge regression decoders for the data averaged in a 200ms window around the maximum of each decoder. **(C)** Transfer decoders trained on one condition and tested on the other. Significant transfer decoding window (cluster-based permutation analysis) for updates is indicated by the shaded area (p<0.01). **(D)**. Abstraction score: within-condition vs transfer decoding ratio. Pitting within-context decoding accuracy and transfer decoding accuracy against each other over time outlines the trajectory of abstraction. Bar plots: maximum value for each stage of the hierarchy. **(E)** Decoding accuracies split for correct (solid lines) and incorrect trials (dotted lines). Only update accuracy is significantly predictive of trial correctness (cluster-based permutation, p<0.01).

### Neural abstraction across object features

The near-perfect feature selection measured at the update stage indicates that participants accumulate evidence only for the relevant feature in 1D contexts. But does evidence accumulation rely on similar neural representations across color and shape contexts? We tested this possibility by measuring the performance of a **transfer decoder**, trained to decode a target object property in one context and tested by decoding the same object property in the other context (Bernardi et al., 2020) (see **Methods**). Transfer decoding performance, expressed as a fraction of within-context decoding strength, reflects the degree of **abstraction** of the associated neural representation. This **transfer decoding** approach confirmed that the aspect (color and shape) of objects is associated with dissimilar neural representations (i.e., no significant transfer decoding from color to shape and vice versa; **Figure 2C**). Interestingly, the evidence provided by objects did not show any sign of transfer decoding across 1D contexts. By contrast, we obtained significant transfer decoding for the updates triggered by objects across color and shape contexts (cluster-level decodability: *p* < 0.01; **Figure 2C**). At this processing stage, the performance of transfer decoding (reflecting the degree of abstraction of the associated neural representation) increased over time, from 46% at 310 ms following object presentation (peak of within-context decodability) to 86% at 800 ms (**Figure 2D**).

### Noise-induced neural selection and abstraction

The decoding results obtained so far indicate that participants accumulate evidence only for the relevant feature (**selection**) using highly similar neural representations of updates in accumulated evidence across color and shape contexts (**abstraction**). We reasoned that this reuse of a single evidence accumulation process across 1D contexts may arise from resource constraints. The existence of resource constraints on evidence accumulation is supported by internal noise in this process – measured from behavior (**Figure S2**). We confirmed the existence of resource constraints from MEG by comparing the decodability of context-relevant updates (which reflect evidence accumulation) preceding correct decisions and errors (**Figure 2E**). We found that decision errors were preceded by reduced decodability of updates (cluster-level difference: *p* < 0.01), but no difference in decodability at earlier object processing stages.

To assess whether resource constraints promote neural selection and abstraction, we trained recurrent neural networks (RNNs) (Yang and Wang, 2020) to perform context-dependent 1D decisions in the presence of internal noise in their recurrent updates (Findling and Wyart, 2024) (**Figure 3A**; see **Methods**). In the absence of internal noise (i.e., no resource constraints), trained RNNs achieved optimal categorization accuracy (**Figure 3B**). As expected, injecting internal noise of standard deviation *σ* in recurrent updates decreased training accuracy (**Figure 3B**), but it simultaneously reduced the dimensionality of RNN activation patterns (participation ratio: 3.9 for *σ* = 0, 2.1 for *σ* = 2; **Figure 3C**; see **Methods**). To assess whether this noise-induced reduction of dimensionality is associated with neural selection and abstraction, we decoded the accumulated evidence from recurrent activation patterns and compared decodability: 1. between the relevant and the irrelevant features (selection), and 2. between within-context decoding and between-context (i.e., transfer) decoding (abstraction; **Figure 3D**). Trained RNNs without internal noise showed no clear sign of selection (6%) nor abstraction (16%; **Figure 3E**). This was true with or without gating the context-relevant feature at the input of the recurrent layer. In the presence of internal noise at levels compatible with human categorization accuracy, trained RNNs acquired both selection (maximum: 54%) and abstraction (maximum: 55%; **Figure 3E**). This finding supports the idea that resource constraints on evidence accumulation promote neural selection and abstraction.

**Figure 3.**
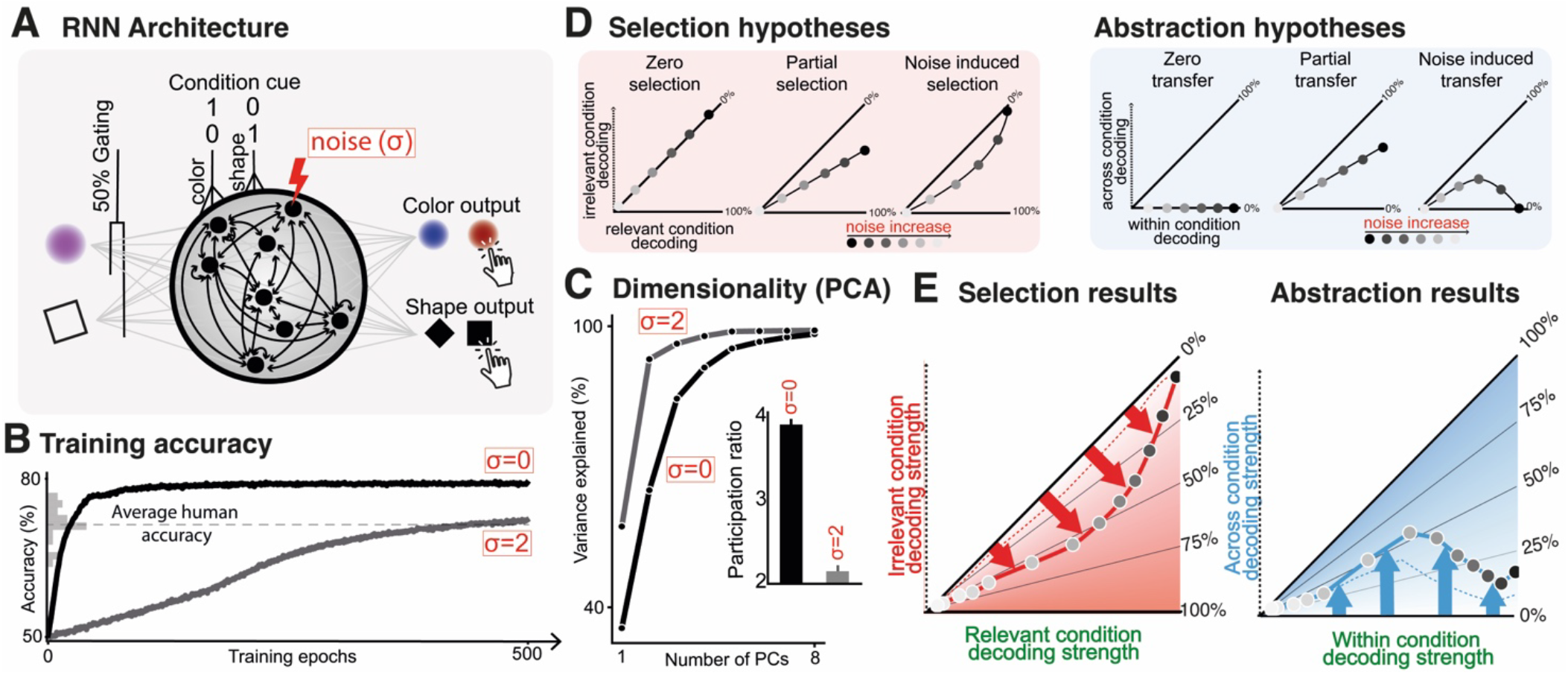
Noise-induced selection and abstraction in RNNs. **(A)** RNNs Architecture: networks received two stimulus inputs (color and shape) and two context inputs. RNNs were trained under varying levels of internal noise in the hidden layer and with 50% gating applied to the irrelevant input (dashed line in panel E show replication without gating - see Methods). The networks were trained to produce accurate categorical decisions for each stimulus sequence. **(B)** Training accuracy over time for noise-free RNNs and RNNs with a moderate level of internal noise (σ = 2). Barplot on the Y axis represents the distribution of human accuracy. **(C)** Dimensionality analysis of noise-free and noisy RNNs. Comparison of the percent of variance explained by the different components of a PCA on the hidden layer activity. Barplot: comparison ot the participation ratio. **(D)** Selection is assessed by decoding the accumulated evidence of both relevant and irrelevant stimulus features from latent space, across different levels of hidden-layer noise. Decoding accuracies were plotted in a two-dimensional space (relevant vs. irrelevant feature decoding). In the absence of selection, both features would be decoded with similar accuracy. In contrast, selective processing is characterized by higher decoding accuracy for the relevant feature than for the irrelevant feature. To assess abstraction, we compared within-condition decoding with cross-condition transfer decoding. **(E)**. Noise-free networks failed to selectively prioritize the relevant feature (no selection) and only moderate sign of abstraction. Adding noise to hidden-layer activity led to robust selection of the task-relevant feature and abstraction or the internal representation. Dashed lines represent the sale analysis in the absence of input gating.

### Neural combination of object features

Analyses of behavior have shown that participants could perform 2D decisions only slightly worse than equally difficult 1D decisions (**Figure 1CD**), and that they do so by combining object features on individual trials (**Figure 1FL**). But **how** does the brain combine object features in 2D contexts? Do resource constraints also explain how the brain performs multidimensional perceptual decisions? To answer these questions, we decoded the same three properties associated with object processing (aspect, evidence, update) under two competing accounts of object processing in 2D contexts, associated with distinct neural representations but identical behavior (**Figure 4**). The ‘segregated processing’ account (**H**_**1**_) posits that object features are accumulated separately and further combined downstream from evidence accumulation (**Figure 4A**, top), whereas the ‘integrated processing’ account (**H**_**2**_) assumes that object features are combined into category evidence upstream from its accumulation (**Figure 4A**, bottom). Geometrically, **H**_**2**_ means that objects are projected on the categorization axis associated with the active context (e.g., from ‘blue square’ to ‘red diamond’). Under **H**_**1**_, neural decodability should be higher for separate features and lower for combinations of features (**Figure 4B**). Under **H**_**2**_, neural decodability should be higher for the relevant combination of features, lower for separate features, and lowest for the irrelevant combination of features (**Figure 4B**). Decoding object properties from spatial patterns of MEG activity under these two accounts of 2D decisions revealed that the processing of aspect and evidence is compatible with **H**_**1**_ (cluster-level difference: all *p* < 0.01; **Figure 4D**), whereas the processing of updates is compatible with **H**_**2**_ (cluster-level difference: all *p* < 0.05; **Figure 4D**). This pattern of findings indicates that the brain initially represents object features separately before collapsing them on a single axis to accumulate evidence only for the relevant combination of features.

**Figure 4.**
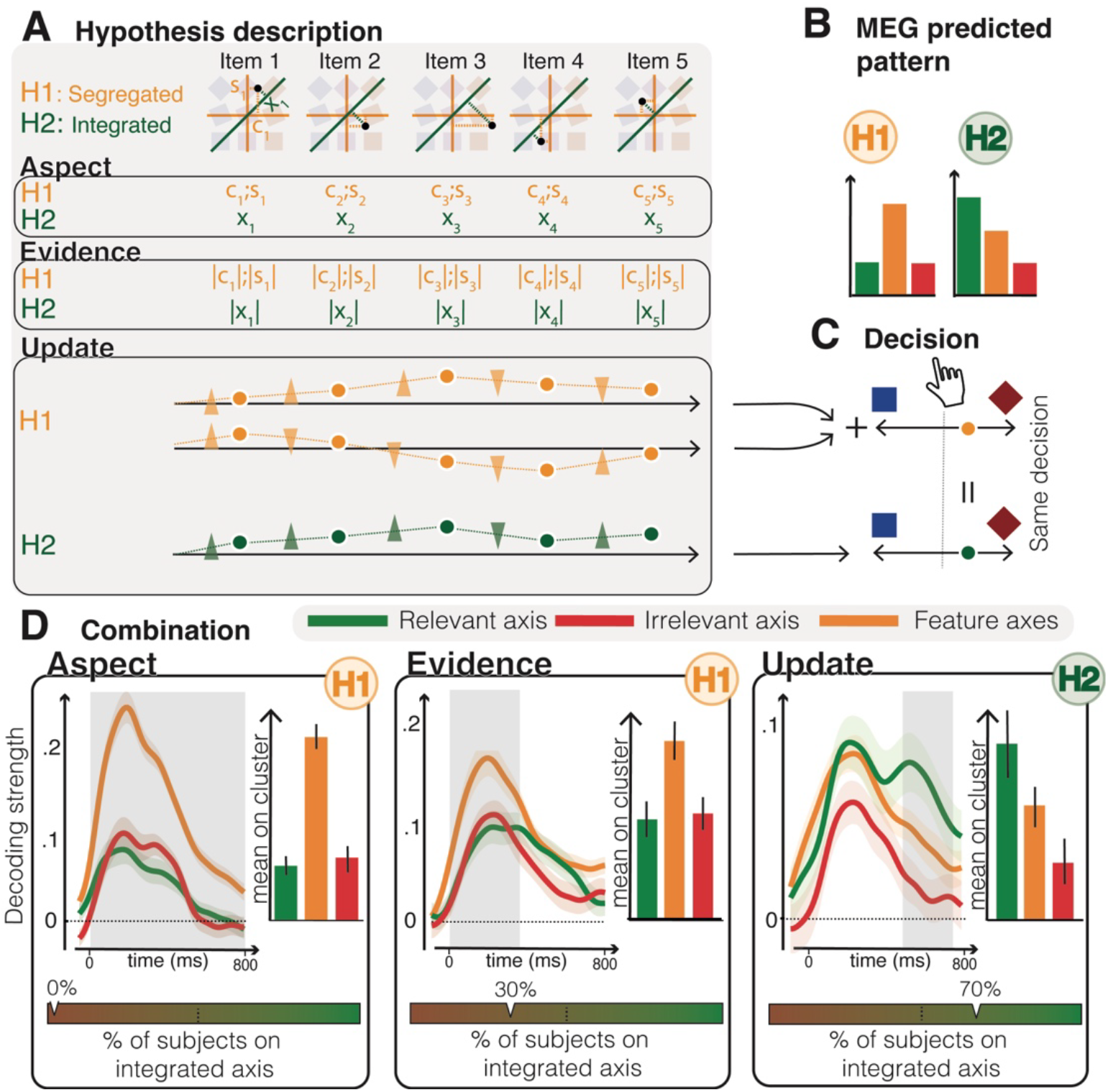
Neural combination of object features. **(A)** Description of the two hypotheses tested at each stage: 1/ *segregated processing* of object features (H1), and 2/ *integrated processing* of the relevant combination of object features associated (H2). The time course represents the total accumulated evidence under the two hypotheses. **(B)** Under H1, neural decodability should be higher for individual object features, and lower for relevant and irrelevant combinations of object features. Under H2, neural decodability should be higher for the relevant combination of object features, lower for individual object features, and lowest for the irrelevant combination of object features. **(C)**. Despite different internal intermediary states, both segregated or integrated updates predict the same behavioral outcome **(D)** Decoding accuracies revealed that the perceptual steps are most compatible with a segregated processing (higher decodability for individual features for aspect and evidence, cluster-level difference: p < 0.01), while the update is preferentially performed over the combination of features (higher decodability for the relevant combination of features, cluster-level difference with individual features: p < 0.05, cluster-level difference with the irrelevant combination of features: p < 0.05.). All these analyses have been replicated by decoding the amount of accumulated evidence from the same data (**Figure S5**).

### NOISE-resilient neural combination

The ‘integrated processing’ account of evidence accumulation in 2D contexts supports the existence of resource constraints that favor the accumulation of evidence for the relevant combination of features over the separate accumulation of color and shape evidence. Consistent with this view, decision errors were preceded by reduced decodability of updates for the relevant combination of features (cluster-level difference: *p* < 0.01), but not for separate features (cluster-level interaction: *p* < 0.05). But does this ‘integrated processing’ account provide objective benefits over the ‘segregated processing’ account in the presence of internal noise in evidence accumulation? To answer this question, we implemented these two accounts in RNNs trained to perform 2D decisions (**Figure 5A**). Segregated RNNs consisted of distinct color- and shape-selective units, whereas integrated RNNs consisted of mixed-selective units (see **Methods**). By decoding the accumulated evidence from recurrent activation patterns of trained RNNs under **H**_**1**_ and **H**_**2**_, we verified that segregated RNNs accumulate color and shape evidence separately (**H**_**1**_), whereas integrated RNNs accumulate evidence for the relevant combination of color and shape (**H**_**2**_; **Figure 5B**). In the absence of internal noise (i.e., under no resource constraints), both types of RNNs achieved optimal categorization accuracy (**Figure 5C**). By contrast, in the presence of internal noise at levels compatible with human categorization accuracy (Findling and Wyart, 2024), integrated RNNs achieved substantially higher accuracy than segregated RNNs with the same amount of training (segregated: 58%, integrated: 69%; **Figure 5CD**). The increased resilience of integrated RNNs to noise was specific to internal noise in recurrent updates: both types of RNNs achieved similar accuracy in the presence of input noise (**Figure 5E**), output noise (**Figure 5F**), or lesion in the hidden layer (**Figure 5G**). The increased resilience of integrated RNNs to noise was also specific to 2D decisions: both types of RNNs achieved similar accuracy when trained to perform 1D (color or shape) decisions (**Figure S6**).

**Figure 5.**
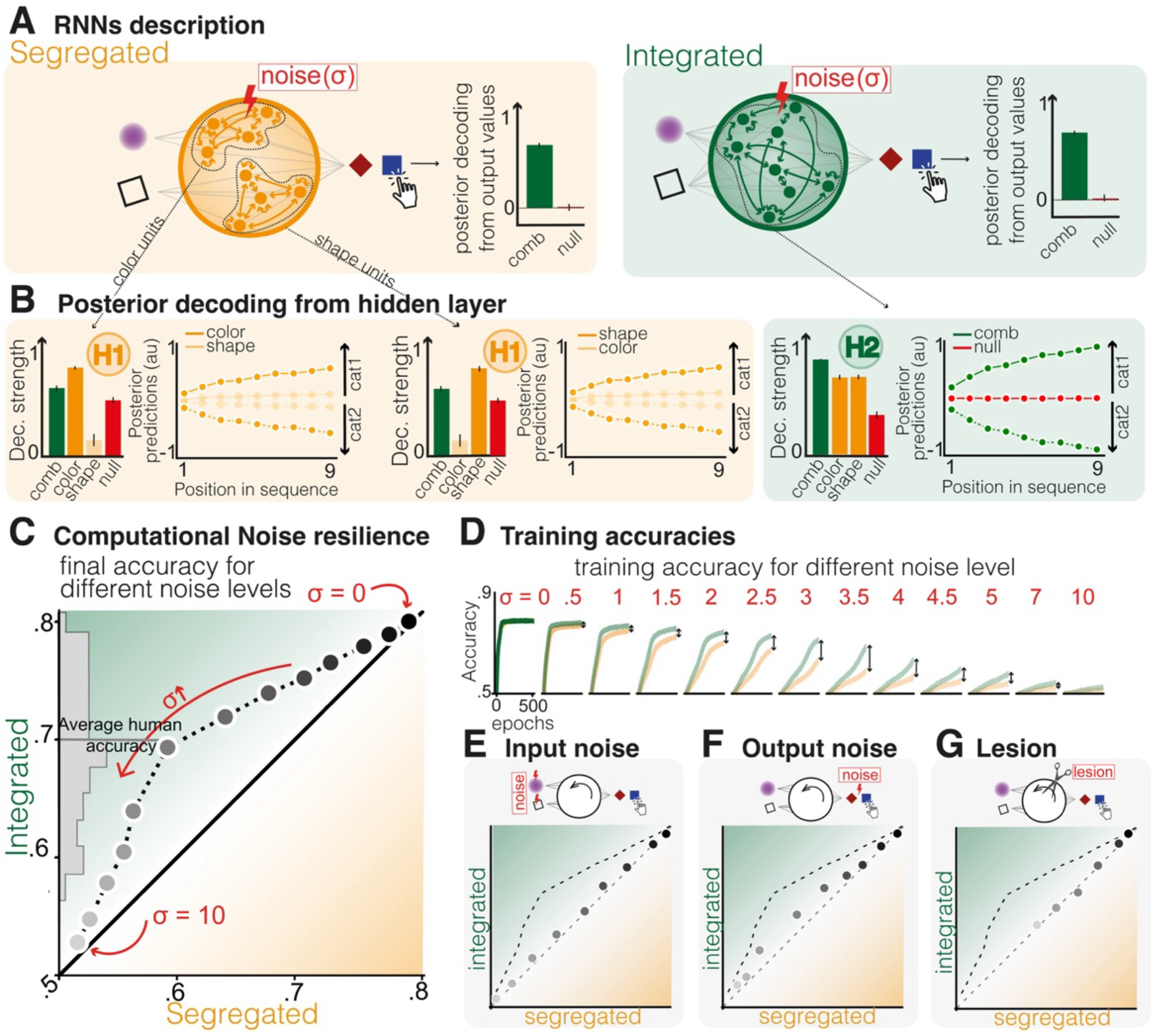
Noise-resilient neural combination. **(A)** Variations of RNNs implementing a different strategy. To ensure segregated accumulation of evidence in split RNN, the hidden layer is split in two independent subparts each receiving a single feature as input. We compared it to a sparse RNN with same number of nodes and connections in the hidden layer (random 50% of its recurrent connections being removed to match split networks’ sparsity). **(B)** Posterior value decoding from the hidden layer: split RNNs implement segregated accumulation while the sparse RNNs implement feature combination. **(C)** In the absence of noise both structure reach optimal performances. Adding noisy recurrent connections impaired more severely single feature split networks compared to multi features (sparse) networks. **(D)** This effect was not due to difference in learning speed between the two types of networks but rather the accuracy convergence they can reach. **(EFG)** RNN sensitivity to input noise **(E)**, output noise **(F)** or hidden layer lesion **(G)**. None of them had a similar distinctive effect between the two types of networks.

### Neural abstraction across 1D and 2D decisions

Combining features prior to their accumulation effectively reduces 2D decisions to a 1D evidence accumulation process (**Figure 4A**). This opens the possibility for the brain to reuse similar neural representations to accumulate evidence not only across color and shape decisions (**Figure 2D**), but even across 1D and 2D decisions. To test this extended form of neural abstraction, we trained a transfer decoder to decode updates in one context (color, shape, mixed 1, or mixed 2) and tested the performance of the decoder by decoding updates in all other contexts (see **Methods**). The resulting **transfer decoding matrix** (**Figure 6A**) showed significant transfer decoding across all context pairs (all Bonferroni-corrected *p* < 0.01). Importantly, a transfer decoder trained to decode updates in 1D contexts provided further support for the ‘integrated processing’ account (**H**_**2**_) over the ‘segregated processing’ account (**H**_**1**_) when tested to decode updates under these two accounts in 2D contexts. Indeed, transfer decoding performance was maximal for the relevant combination of features, lower for separate features, and lowest for the irrelevant combination of features (cluster-level difference: all *p* < 0.01; **Figure 6B**). To examine and compare the dynamics of associated neural representations across contexts, we trained a final **abstract decoder** to decode updates from the same spatial patterns of MEG activity across all 1D (color, shape) and 2D (mixed) contexts. The identified neural representation was highly dynamic in time: abstract updates were best decoded by orthogonal subspaces of MEG activity spread by only 300 ms (**Figure 6C**; see **Methods**). Plotting the trajectory of decodability projected on subspaces trained at 250 ms and 550 ms (**Figure 6D**) revealed highly similar trajectories across 1D and 2D contexts, consistent with the idea that the human brain reuses the same neural representation to accumulate evidence across 1D and 2D decisions.

**Figure 6.**
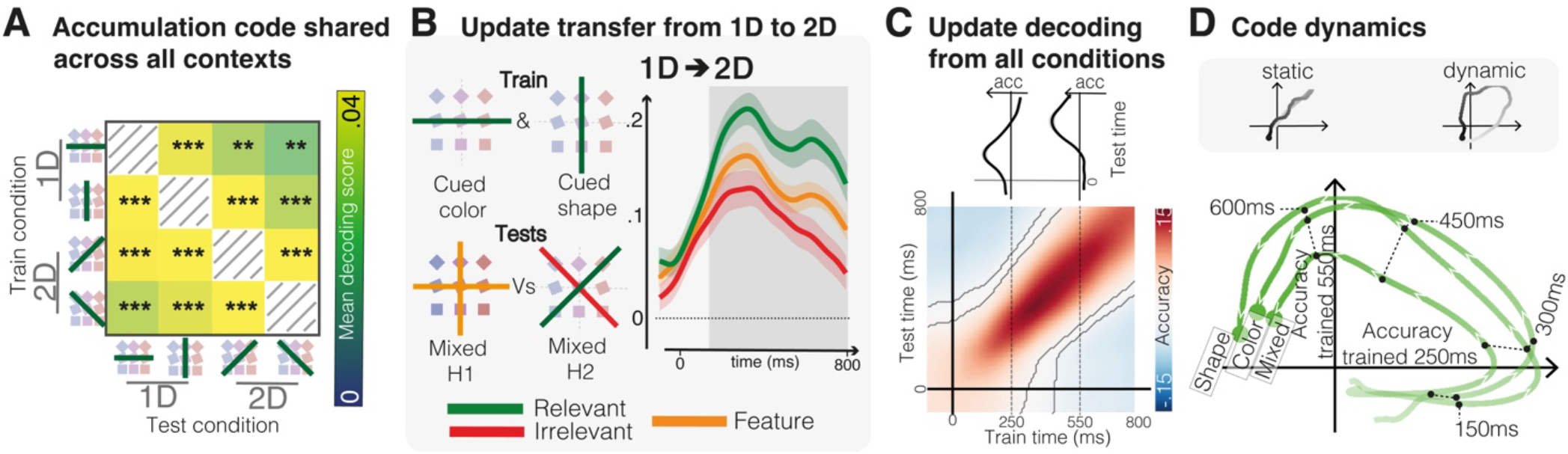
Neural abstraction across 1D and 2D contexts. **(A)** We tested neural abstraction between 1D and 2D conditions by training the update decoder on MEG data from any of the four conditions (Color, Shape, Mixed1 or Mixed2) and tested on data from any other conditions. The average accuracy (fromm 0 to 800 ms was significantly above chance between every pair of conditions (all ps<0.01, Bonferroni corrected). **(B)** update decoder trained on the 1D conditions (Shape & Color) and tested on the mixed conditions, for the individual features, the relevant and the irrelevant combination of features replicated both the preference of humans for update of the combination of features and the abstraction of the update code between 1D and 2D conditions (clusters ps<0.001). **(C)** We computed the Generalization Across Time (GAT) matrix of an update decoder trained simultaneously on all conditions. The resulting matrix displayed a diagonal pattern, indicative of a dynamic coding scheme (in contrast, a static code would produce a square-shaped GAT pattern). We selected two training time points (250 ms and 550 ms) and plotted the decoding accuracy of each corresponding decoder across all testing times. **(D)** To characterize the temporal dynamics of the code, we plotted the decoding accuracies of the two decoders (trained at 250 ms and 550 ms) against each other. A static representation would yield highly correlated accuracies across time, whereas a dynamic code would generate a circular trajectory. We performed this analysis separately for the Shape, Color and Mixed, conditions, and observed a similar dynamic code for the update across all conditions.

## Discussion

### Decision flexibility beyond feature selection

We have shown that humans can not only select but also combine multiple object features (both within and across sensory modalities) and that the brain supports this flexibility by reusing a shared, abstract and dynamic neural subspace for accumulating evidence – rather than relying on separate subspaces for individual object features. This finding extends the definition of decision flexibility to multidimensional perceptual decisions. Indeed, existing work typically equates decision flexibility with the context-dependent **selection** of a decision-relevant sensory feature, and has therefore aimed at identifying neural computations that support this flexible feature selection process (Mante et al., 2013; Okazawa and Kiani, 2023; Pagan et al., 2025). Here we have extended the definition of decision flexibility by studying the **combination** of multiple sensory features required to perform multidimensional decisions – a class of decisions that is common in everyday perception, but rarely studied in the research laboratory. For this purpose, we have developed a multidimensional categorization task – instantiated in the visual domain (**Exp. 1**) and in the audio-visual domain (**Exp. 2**) – where tested participants need to select but also combine multiple features of presented objects to make accurate decisions. We found that humans can combine multiple features, even across sensory modalities, to make accurate decisions. In the visual domain, where we recorded MEG data while participants made 2D decisions, we found that both selection and combination are supported by the neural **abstraction** of evidence accumulation – the reuse of shared neural representations to accumulate evidence collapsed onto the category axis across 1D (selection-based) and 2D (combination-based) perceptual decisions. These findings shed new light on the role of abstraction in flexible perceptual decisions.

### Evidence accumulation as a cognitive bottleneck

Several observations point toward the existence of a cognitive bottleneck (i.e., strong resource constraints) at the evidence accumulation stage. First, consistent with previous work (Drugowitsch et al., 2016; Wyart and Koechlin, 2016; Findling and Wyart, 2021), internal noise in evidence accumulation accounted for the majority of participants’ decision errors across all contexts. In line with this, moment-to-moment fluctuations in the decodability of updates (reflecting evidence accumulation) from MEG activity predicted the accuracy of the resulting decision. Second, neural selection increased across processing stages, reaching its maximum at the update (evidence accumulation) stage, where resource constraints are therefore likely to be the strongest. This observation is consistent with several findings in both human and non-human studies (Pestilli et al., 2011; Mante et al., 2013; Siegel et al., 2015; Brincat et al., 2018; Birman and Gardner, 2019; Barbosa et al., 2023; Zhang et al., 2025; Pagan et al., 2025). It is also consistent with an additional condition in which the 1D context (color or shape) could change from one trial to the next and was revealed at response time (**Figure S7**). This condition required participants to accumulate color and shape evidence in parallel, and select the relevant sensory feature at response time. We obtained clear signs of neural **competition** in the decodability of color and shape updates from MEG activity in this condition – in line with recent work (Kang et al., 2021). Third, the brain reduced 2D decisions to a single evidence accumulation process by accumulating evidence for the relevant combination of features, rather than accumulating color and shape evidence separately. Importantly, this conclusion held true when decoding the magnitude of accumulated evidence (instead of updates) from spatial patterns of MEG activity (**Figure S5**).

### Decision flexibility through neural abstraction Transfer decoding

(also named CCGP for ‘cross-condition generalization performance’ in previous work) revealed that the brain not only reduces 2D decisions to a 1D evidence accumulation process, but also reuses the same neural representation to accumulate evidence about color, shape, and combinations of these two sensory features. This finding means that evidence accumulation can operate in an abstract representational space that is decoupled from upstream sensory features (Benjamin et al., 2026; Bernardi et al., 2020; Drevet et al., 2024; Flesch et al., 2022; Luyckx et al., 2019; O’Connell et al., 2012; Sheahan et al., 2021). This representational space can also be decoupled from downstream effectors because, in our task, the mapping between categories and effectors was unavailable during evidence accumulation. From this perspective, neural selection and combination correspond to two instances of flexible projections of high-dimensional sensory representations onto a decision axis that is used to accumulate evidence over time. This low-dimensional ‘read-out’ could be implemented by populations of neurons with mixed selectivity in associative prefrontal and parietal regions (Cazettes, 2025; Panichello and Buschman, 2021; Rigotti et al., 2013), which we successfully modeled using RNNs with 2D selectivity to color and shape. In this ‘flexibility through abstraction’ account, cognitive flexibility arises from the tuning of this low-dimensional projection in a context-dependent fashion, rather than from the stacking of context-specific subspaces for evidence accumulation. Future research should not only investigate the fine-grained neural mechanisms that support this flexible low-dimensional projection, but also the other types of projections (beyond neural selection or combination) that humans can apply in the broad range of conditions encountered in everyday life.

### Toward a resource-efficient account of decision flexibility

By training RNNs to perform 1D and 2D decisions in the presence of internal noise in their recurrent updates, we could test whether resource constraints promote the three properties of decision flexibility that we have identified in the human brain: **selection, combination** and **abstraction** (Kanwisher et al., 2023). In the absence of internal noise, RNNs failed to acquire selection and abstraction during 1D decisions, and the combination of features did not provide any objective benefits during 2D decisions. By contrast, when internal noise was introduced at levels consistent with human categorization accuracy, trained RNNs showed clear signatures of selection and abstraction during 1D decisions, and the combination of features upstream from evidence accumulation conferred greater robustness to internal noise during 2D decisions. These results shed new light on the multiple effects of internal noise in decision-making. By nature, random noise in evidence accumulation (recurrent updates in RNNs) corrupts the decision variable and triggers suboptimal decision variability. But internal noise also shapes neural computations in ways that enhance decision flexibility. At fast (inference) time scales, recent work has shown that internal noise promotes decision flexibility under novel sources of uncertainty (Dapello et al., 2020; Findling and Wyart, 2024). At slow (optimization) time scales, our results further suggest that internal noise promotes the acquisition of selection, combination and abstraction during perceptual decision-making. Recent work has shown that an explicit low-rank constraint on the dimensionality of RNN activations (Dubreuil et al., 2022; Mastrogiuseppe and Ostojic, 2018) can also favor the reuse of the same activation subspaces across different contexts of the same task – i.e., abstraction (Beiran et al., 2023). Future research should therefore investigate whether the dimensionality of RNN activations mediates the beneficial effects of internal noise on decision flexibility.

### Abstraction beyond perceptual decisions

The **neural abstraction** process we have identified during multidimensional perceptual decisions is likely not to be specific to perception. Indeed, the same class of computational problems arises whenever behavior requires combining heterogeneous sources of information under a common criterion (e.g., integrating verbal and non-verbal cues when judging another person’s intent, pooling symptoms of different kinds when reaching a medical diagnosis): multidimensional input must be combined and accumulated to deliberate toward an accurate decision. Our **‘flexibility through abstraction’** account proposes that the human brain solves this problem not by running separate accumulation processes for each source of evidence, but by using a single evidence accumulation process collapsed on the decision (relevant) axis. This account makes a testable behavioral prediction for future work: individuals relying on more abstract representations should switch between contexts at lower cost, and should generalize more readily to feature combinations that they have never encountered. It also reframes the relationship between capacity limits and flexibility, suggesting that the constraints usually treated as a source of human suboptimality are also what make context-general strategies worth acquiring.

Together, these findings highlight abstraction, the ability to form latent representations that generalize across diverse situations, as a key, yet understudied component of decision flexibility. By examining both **how** humans make flexible perceptual decisions – by projecting relevant sensory features onto a shared, abstract evidence axis prior to its accumulation – and **why** humans adopt this cognitive strategy – due to resource constraints on evidence accumulation – we outline a comprehensive framework in which resource constraints simultaneously degrade decision accuracy while promoting cognitive strategies that enhance decision flexibility.

## Methods

### Participants

#### Online participants (Exp. 1-2)

Two independent samples were recruited online through the Prolific platform. In the visuo-visual experiment (**Exp. 1**), *N* = 121 adult participants completed the task (63 females). In the audio-visual experiment (**Exp. 2**), *N* = 131 adult participants completed the task (68 females); all of them passed a headphone screening test administered before the task, and participants who failed this screening test did not proceed to the main experiment. No participant took part in both experiments. All participants gave informed consent before starting and were compensated £9.50 per hour for their time. The study was approved by the Inserm Ethical Review Committee (IRB00003888 on 13/11/2018).

#### MEG participants (Exp. 3)

24 French adult participants (14 females, 18-30 years old, all right-handed) participated in the study. All participants had normal hearing, normal or corrected-to-normal vision, and no history of neurological nor psychiatric disorders and were not color-blind. All tested participants gave a written informed consent before taking part in the study, which received ethical approval from relevant authorities (Comité de Protection des Personnes Ile-de-France VI, ID RCB: 2017-A01778-45). Each participant received a monetary compensation of 80 € for their participation in the study.

### Experimental procedures

#### Stimuli

Objects (**Figure 1**) were colored shapes (**Exp. 1 & 3**), or beeping color flashes objects (**Exp. 2**). Color is varying from blue to red (**Exp. 1, 2 & 3**), orientation varying from 0 to 45 degrees (**Exp. 1 & 3**) and pitch varying from 220 Hz (A3) to 1760 Hz (A6) (**Exp. 2**). They were presented sequentially at fixation. For the MEG experiment (**Exp. 3**) we controlled distance to the screen so that the size of the objects was 2.5 degrees of visual angle.

#### Online task design (Exp. 1-2)

Participants were presented with a sequence of objects and were asked to make a 2-AFC judgment about the generative category of the sequence, by accumulating successive information contained in each item of the sequence. Sequences were composed of 4, 8, 12, or 16 consecutive stimuli presented at 4 objects/sec. After the presentation of the sequence, participants were presented visually with the two categories of interest on the left and right of fixation, and had to report to which of the two categories the sequences adhered the most by pressing a button. We presented 4 different conditions to participants: **Feature 1** (color), **Feature 2** (shape in **Exp. 1**, auditory pitch in **Exp. 2**), **Mixed 1** and **Mixed 2**. Participants were informed about the categorical dimension of interest before the sequence presentation (see **Figure 1BH**). In the **Color** condition, they had to categorize the sequence over the color axis (blue or red) while ignoring the other feature. In the **Shape** condition (**Exp. 1**), participants had to judge over the orientation dimension (square: 0 deg orientation, or diamond: 45 deg orientation) while ignoring the color. In the **Pitch** condition (**Exp. 2**), participants had to judge over the pitch dimension (low: 220 Hz, or high: 1760 Hz) while ignoring the color. In **Mixed 1** and **Mixed 2** conditions, they had to categorize over two-dimensional axes (e.g., ‘blue square’ or ‘red diamond’ in the Mixed 1 condition for Exp. 1, or ‘high-pitched blue’ or ‘low-pitched red’ in the Mixed1 condition for Exp. 2).

#### MEG task design (Exp. 3)

This task is an MEG-compatible variant of **Exp. 1**. Sequences were composed of 2, 4, 6, or 8 consecutive stimuli presented at 2 or 4 Hz. For all analyses, trials at 2 and 4 objects/sec were aggregated. Stimuli were flashed on the screen for 167 ms and the ISI was 167 ms for 4 objects/sec presentation and 333 ms for 2 objects/sec presentation. After the presentation of the sequence, participants were presented with the two categories of interest and had to report to which of the two categories the sequences adhered the most by pressing a button. We presented 5 different conditions to participants. For the four cued conditions (**Shape, Color, Mixed1** and **Mixed2**), participants were informed about the categorical dimension of interest before the sequence presentation (see **Figure 1B**). In the **Shape** condition, participants had to judge over the orientation dimension (square: 0 deg orientation, or diamond: 45 deg orientation) while ignoring the color. In the **Color** condition, they had to categorize the sequence over the color axis (blue or red) while ignoring the orientation of the stim. In **Mixed1** and **Mixed2** conditions, they had to categorize over two-dimensional axes (blue square or red diamond in the **Mixed1** condition, blue diamond or red square in the **Mixed2** condition). An additional condition was added in which participants were not informed about the relevant dimension (color or shape) before the presentation of the sequence but only at the end of the sequence when prompted for a response. Sequences were presented in mini-blocks of 16 trials of the same condition (**Figure 1B & Figure S7**).

#### MEG recording procedures

Each participant was fitted with electrocardiogram (ECG) and electrooculogram (EOG) electrodes, as well as Head Position Indicator (HPI) coils to track their head position in the MEG scanner helmet throughout the experiment. The perceptual decision task was explained to the participant, followed by a training session prior to MEG recordings. Participants were instructed to refrain from blinks and head movements in the middle of sequences of stimuli. Participants underwent 8 MEG recording runs, with self-paced breaks between each recording run.

### Behavioral analyses

#### Rejection

For online behavioral experiments (**Exp. 1-2**), participants were excluded from all analyses if their categorization accuracy in the 1D contexts, pooled across the two contexts, was not significantly above chance (one-tailed binomial test, *p* < 0.05), or if their accuracy was at or below chance (chance: 50% correct) in either 1D context. This criterion removed 9 of all 121 participants in the visuo-visual experiment (**Exp. 1**) and 25 of all 131 in the audio-visual experiment (**Exp. 2**).

#### Performance

For each participant and each condition, we computed the percentage of correct categorization of sequences compared to the generative category of the sequence. For **Mixed** contexts, we computed one-sample *t*-tests to assess significance of the performance above chance level (**Figure 1CI**; chance; 50% correct, *p* < 0.001).

#### Psychometric analyses

For each participant and each condition, we estimated the sensitivity to the decision variable corresponding to the amount of Bayesian evidence (log-posterior ratio) separately for the active decision axis (e.g., color in the **Shape** condition), and for the null decision axis (e.g., shape in the **Color** condition) through logistic regression, using a logit link function (**Figure 1DJ**). We averaged the sensitivity coefficients across participants for each condition, and compared them to zero using one-sample *t*-tests (**Figure 1EK**).

#### Strategy comparison

For each participant, three strategies were fit to the choices made in the **Mixed** contexts. A random strategy ignored the evidence and had a category bias (1 parameter); a 1D strategy read a single feature on each trial, color with probability π and the other feature otherwise, and was therefore a mixture of two unidimensional logistic policies (6 parameters); a 2D strategy that combined both features on every trial (4 parameters). Both decision strategies included a lapse rate. Parameters were estimated by maximum likelihood with a regularization L2 penalty of 0.05 on the feature weights, to prevent fits with extreme values. Strategy prevalence in the population was then estimated by random-effects Bayesian model selection, taking −AIC/2 (Akaike Information Criterion) as log model evidence. Recovery of these three strategies showed that they are correctly differentiable from the data – *p*(simulated|recovered) > 0.70 for all strategies.

### Cognitive computational model fitting

#### 1D evidence accumulation model

To fit the trials from 1D conditions (**Shape** and **Color**), we used a cognitive model of evidence accumulation comprising 4 key parameters: 1/ stimulus bias to either category (*δ*_*acc*_) 2/ accumulation leak (*α*_acc_), 3/ accumulation noise (σ_acc_), and 4/ response noise (σ_rep_). Biased evidence from successive inputs (*x*_*i*_) was accumulated with a temporal leak across successive update steps:

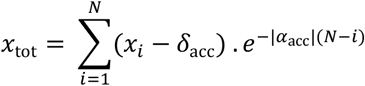

The corresponding accumulated variance is therefore:

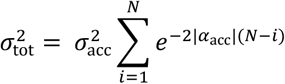

So that the final evidence for a category follows 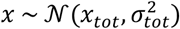. From this final evidence *x*, the probability of the choice is computed through the cumulative normal function with mean *x* and var-iance 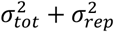, accounting for response selection noise. We also accounted for a probability of flipped responses in which participants used the wrong response mapping (e.g., using the ‘red’ key to report ‘blue’ in the **Color** condition). This probability of flipped responses was fiRed along the other parameters.

#### 2D evidence accumulation model

**Mixed** conditions were fitted with a model with the same parameters as the 1D model, but with an additional weighting factor *ω* controlling for the projection of the 2D color and shape onto a single combination: 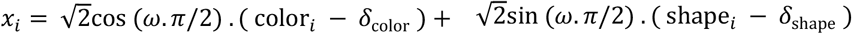 for the **Mixed1** condition or 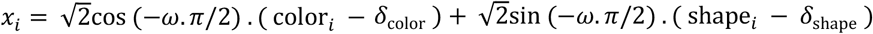 for the **Mixed2** condition. The weighting factor *ω* ranges between 0 (color only) and 1 (shape only), with ½ corresponding to the perfect combination of color and shape accumulation with no bias for any feature. The accumulation and decision processes were identical to the 1D model.

#### Fitting procedures

We implemented these models and fitted them to participants’ choices. Per-trial choice log-likelihoods were estimated and then combined with prior distributions on parameter values (uniform priors across the tested range) to get a logposterior estimate. The log-posterior was fed to the Variational Bayesian Monte Carlo (VBMC) algorithm [Acerbi, 2020; version 1.0; https://github.com/lacerbi/vbmc] returning a variational approximation of the full posterior and a lower bound on the log-marginal likelihood. We considered the posterior mean (and standard deviation) to obtain best-fitting parameter values (and standard deviation).

### MEG preprocessing

#### Data acquisition

MEG data were recorded using a whole-head Elekta Neuromag TRIUX system (Elekta Instrument AB, Stockholm, Sweden) composed of 102 triplets, each comprising one magnetometer and two orthogonal planar gradiometers, at a sampling frequency of 1000 Hz. Prior to the experiment, each participant’s head shape was digitized together with four additional head position indicator (HPI) coils, whose positions were used to monitor and correct for small head movements across blocks. Head position in the scanner was measured before each block.

### Data preprocessing

We first applied signal source separation (SSS, MaxFilter algorithm) to remove ambient noise and realign blocks to the average head position. MEG data were further preprocessed using position. MEG data were further preprocessed using MATLAB and the FieldTrip toolbox. Data were subsampled to 500Hz and epoched between [- 200;1800]ms. Epochs were linearly detrended and band-pass filtered between 0.5 and 8 Hz. For all subsequent steps and analyses, we made the conservative choice of keeping all epochs to avoid unbalancing the design and introducing an arbitrary cutoff threshold. We further ran ICA using the FieldTrip built-in function for each participant and each run separately. Visual inspection of the topographical maps and component time courses of the 20 first components allowed rejections of artifactual components (eye movements, cardiac activity…). Hilbert transform was applied to the MEG data. For all decoding analyses, we used real and imaginary parts at each time point to optimize their accuracies. This enables considering both instantaneous brain activity but also its dynamical change in proximal horizon.

#### Data quality assessment

We assessed individual MEG data quality independently from contrasts of interest, by decoding the aspect on **Color** and **Shape** conditions for each participant. A single participant was excluded from further analysis, whose object aspect (color and shape) could not be decoded better than chance (average decoding score < 0 between - 200 ms to +800 ms following object presentation).

### MEG analyses

#### Definition of regressors

We defined four regressors for the analyses. Let *x*_*i*_ ∈ [−1,1] denote the signed visual input associated with the stimulus presented at position i in the sequence. In the color domain, *x*_*i*_= −1 corresponds to a fully blue stimulus, *x*_*i*_ = +1 to a fully red stimulus, and intermediate values reflect the stimulus’ position along the blue–red continuum. In the shape domain *x*_*i*_ varies from -1 (square) to 1 (diamond). The **aspect** regressor was defined directly as *A*_*i*_ = *x*_i_. The **evidence** regressor quantified the amount of sensory evidence conveyed by the stimulus and was defined as the absolute value of the visual input, *E*_*i*_ = |*A*_*i*_|. The **update** regressor indexed belief updating and was defined as *U*_*i*_ = *x*_*i*_ * *sign*(*B*_*i−1*_) where *B*_*i*−1_denotes the accumulated belief after integrating all stimuli up to position i−1 in the sequence (see **Figure 2A** for a visual description of each regressor). Finally, the **accumulated** (posterior) regressor (**Figure S5**) corresponded to the amount of accumulated belief at time i, recursively defined across the sequence *P*_*i*_ = |*B*_*i*_| . Instead of considering the theoretical values of the regressors described above, we considered the corresponding internal variables of the cognitive model, best fitted to each participant individually, to account for individual differences in participants’ biases and accumulation parameters (preference for a category, recency effect, accumulation noise…). We used particle filtered parameter estimates, filtered on individual response for each trial to match regressor adequation with participants’ brain state during each trial.

#### Neural decoding procedures

For each decoding analysis, we computed ridge regressions (*λ* = 500) with 10-fold cross-validation to predict regressor values at each time point *t*.

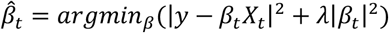

The regressions were performed conjointly on the real and imaginary parts of the Hilbert transformed signal for all magnetometers (we replicated all results from **Figure 2B & Figure 4D** using gradiometers; Figure S8). Once fitted, we could obtain predictions on the test left out fold 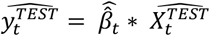. These predictions 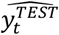 were smoothed with a moving average window of 200ms to improve signal/noise ratio. The obtained smoothed predictions were compared to true values by computing Pearson correlation scores. These correlation scores were Fisher-transformed (to recover normality) and averaged across participants. In **Figure 2A**, the green line (focused axis) corresponds to the average correlation along the shape axis for the **Shape** condition and the color axis for the **Color** condition (focused axis), and the red line to the average correlation of decoding score along the shape axis in the **Color** condition and the color axis in the **Shape** condition (ignored axis). In **Figure 4**, for 2D decisions, the orange line corresponds to the decoding of the values considering segregated processing, while the green line corresponds to the decoding considering integrated processing. The red line corresponds to the decoding over the axis orthogonal to the decision category. This axis shares similar collinearity with the segregated (color and shape) axes as the integrated axis but is irrelevant to the decision. Any decision vs orthogonal axis difference in the decoding is thus to be interpreted as decision relevant computation rather than collinearity with color and shape axes. We further checked that the significance of main results was robust to different smoothing windows of the raw predictions of the decoders. We replicated all results from **Figure 2A & Figure 4B** with other smoothing values; all of which yielded similarly significant results (from no smoothing to 400-ms smoothing).

#### Statistical procedures

To assess significance of each decoding time-course above zero, we performed a cluster-based permutation method across participants (implemented with MNE, 10,000 permutations, threshold=0.05) in the [0-800]ms time window. P-values reported in the text correspond to the significance of each cluster. Significance between two decoding scores was assessed using the paired cluster-based permutation method. For the specific correct vs incorrect contrasts (**Figure 2B** / **Figure S4B** / **Figure S7B**), we used one-tailed tests reflecting the directional hypothesis that correct trials lead to better encoding than incorrect ones.

#### Temporal generalization procedures

For the Generalization Across Time matrix (GAT; Figure S4A) we used exactly the same approach, but each decoder trained at a given time *t*_1_ was further used to predict every other time point 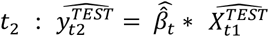. We further used the same z-scored smoothed Pearson correlation as accuracy metric and assessed statistical significance using the 2D temporo-temporal cluster-based analysis.

#### Cortical source localization

MEG cortical source reconstruction was performed using MNE-Python on epoched magnetometer data, using the fsaverage template anatomy. Noise covariance matrices were estimated from the pre-stimulus baseline using a cortically constrained source space (oct6) and a single-layer BEM was computed on fsaverage, and forward solutions were generated using magnetometer sensors only. Source estimates were computed for each individual trial using an L2-regularized solution using MNE function make_inverse_operator with λ2 = 0.05.

#### Neural decoding in the cortical source space

To illustrate the source estimate of each stage of the neural hierarchy, we reproduced the decoding of each code (aspect – evidence – update), on the focused axis only – decoding along the shape axis for the **Shape** condition, and on the color axis for the **Color** condition. Source-space decoding was performed using a searchlight-based ridge regression approach. We computed a single source space decoding of the signal average of a [-100, +100]ms time window around the maximum of the sensor space time resolved decoder. We used a searchlight approach with a geodesic distance threshold (2 cm).

Like in the sensor space, we used a ridge regression (*λ* = 500) with 10-fold cross-validation to predict regressor values. Accuracy was further computed as the Pearson correlation between predicted and true regressor values. **Figure 2A** displays the average correlation score of all participants. To prevent double dipping with sensor space decoding, we made no statistical analyses of source decoding accuracies as these have been computed around sensor space decoding maximum. These maps are presented for visualization of the most probable location of the already demonstrated decoding accuracies.

### Modeling feature selection and abstraction in RNNs

#### Network structure

We employed custom recurrent neural networks (RNNs) implemented with PyTorch. The model consists of a single 32-unit recurrent hidden layer with additive input and recurrent contributions, followed by two parallel output heads. The network takes as input a sequence of both features (color and shape) as well as a two-dimensional one-hot encoding of the relevant condition for this trial (**Color** or **Shape**). For each element of the sequence, the network updates its hidden state (*h*_i_) by combining the current input (*i*_*i*_) with the previous hidden state (*h*_i−1_), hidden with linear transformations (weight matrix input to hidden ***W***_*ih*_, weight matrix hidden to hidden ***W***_*hh*_ and a bias) and applying hyperbolic tangent non-linearity:

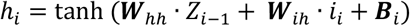

Networks were trained on a similar task as humans, to make categorical judgments about the sequence of input. The network produces two output vectors, each constrained to the interval [0,1] by a sigmoid nonlinearity:

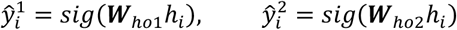

Only the last outputs of the sequences are considered to supervise the learning. We computed a loss for each head independently using the binary crossentropy (BCE), color for the first output and shape for the second output. The total loss was the sum of the **color** head loss for the Color trials and the shape head loss during the **Shape** trials:

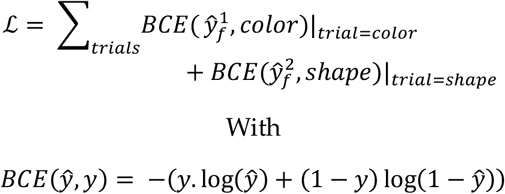

We modeled noisy accumulation of evidence by injecting zero-mean Gaussian noise of variance σ to the hidden pre-activation during each time step 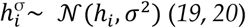. When σ=0, the network reduces to a deterministic RNN. We also implemented two gating by multiplying the irrelevant feature of the input by 0.5.

#### Network training

Hidden states were initialized to zero at the start of each sequence. Networks were trained on sequences corresponding to **Color** and **Shape** conditions. Training batches were composed of sequences of varying lengths (from 2 to 9 items). For each sequence length, an equal number of sequences was generated per batch. Category labels were balanced across conditions and shuffled within each batch. We also implemented gating in which task-irrelevant input dimensions were multiplicatively attenuated by a fixed selection ratio (0.5), simulating selective gating of sensory information at earlier stages (aspect and evidence). This manipulation was applied symmetrically across conditions. The network parameters were optimized using the Adam optimizer with a learning rate of *lr* = 10^−3^. To assess the effect of internal noise on learning and performance, additive Gaussian noise was injected into the hidden-state dynamics during training. Multiple noise amplitudes were tested, ranging from deterministic (σ=0) to strongly stochastic regimes. For each noise level, an ensemble of independently initialized and trained models (n=100) was considered. All models shared the same architecture and training procedure, differing only in noise amplitude.

#### Network testing

After training, models were evaluated on new sequences (not used during training) of lengths 3, 6 or 9. During those evaluation sequences, we decoded at each time step (presentation of each stimulus as input), from the hidden space, the posterior belief accumulated through the sequence for both color and shape dimensions using a similar logistic regression with *λ* = 500 as we used for human MEG decoding. We also estimated abstraction of the posterior belief representation by training a decoder on the **Color** condition and testing it on the **Shape** condition (and vice-versa). For each noise level and gating factor, we plotted the decoding accuracy of the relevant condition against the irrelevant one to assess selection. A similar decoding accuracy would reveal no selection while a lower accuracy for the irrelevant condition is indicative of a selection process in the RNN (**Figure 3DE**). Similarly, we plotted the cross-decoding across conditions against the decoding of the relevant axis to estimate abstraction. No abstraction would be revealed by null cross decoding accuracy, while full abstraction would be revealed by equivalent cross-decoding and relevant feature decoding accuracies (**Figure 3DE**).

### Modeling feature combination in RNNs

#### Network structure

We trained two types of identical RNNs in the 2D (**Mixed**) conditions, only differing in their internal architecture (**Figure 5A**): in one network, the hidden layer was split into two independent subspaces:

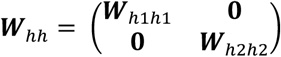

each receiving input from a single feature of the stimulus, whereas in the other, the hidden layer was not split:

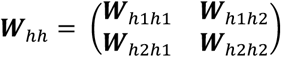

To match the number of trainable parameters in the hidden layer, ***W***_*hh*_ was sparsified with a random 50% of its weights fixed to 0. Similar to RNNs used for selection and abstraction investigation, we used a 32-units recurrent hidden layer with additive input and recurrent contributions.

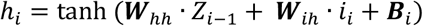

We modeled noisy accumulation of evidence by injecting zero-mean Gaussian noise of variance σ to the hidden pre-activation during each time step.

#### Network training

Networks were trained on sequences corresponding to a **Mixed** condition. Training batches were composed of sequences of varying lengths, with sequence lengths varying from 2 to 9. For each sequence length, an equal number of sequences was generated per batch. Category labels were balanced across conditions and shuffled within each batch. The network parameters were optimized using the Adam optimizer with a learning rate *lr* = 10^−3^. To assess the effect of internal noise on learning and performance, additive Gaussian noise was injected into the hidden-state dynamics during training. Multiple noise amplitudes were tested, ranging from deterministic (σ=0) to strongly stochastic regimes. For each noise level, an ensemble of independently initialized models (n=100) was trained.

#### Network testing

After training, models were evaluated on new sequences (not used during training) of lengths 3, 6 or 9. We computed accuracy for each type of network and plotted split vs sparse network accuracies as a function of hidden layer noise levels (**Figure 5C**).

#### Controls

We reproduced exactly the same procedure with noise corrupting the input (**Figure 5E**):

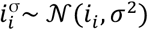

or the output (**Figure 5F**):

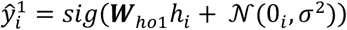

or by introducing lesions to the hidden layer, by clamping a fraction of connection weights to zero after training) (**Figure 5G**).

## Acknowledgments

We are grateful to Hugo Trad for data collection as well as all participants who took part in this study. We thank Maxime Maheu, Mathias Sablé-Meyer and Christopher Summerfield for input and discussions on a previous version of this work.

## Authors contributions

Conceptualization: LB, VW; Methodology: LB, VW; Investigation: LB, JD, VW; Visualization: LB, VW; Funding acquisition: VW; Project administration: VW; Supervision: VW, Writing – original draft: LB, VW; Writing – review & editing: LB, JD, VW

## Code and data availability statement

All preprocessed data along with analyses codes and simulation codes can be found at the following address: https://osf.io/7exkd/

## Supplementary information for

**Figure S1:**
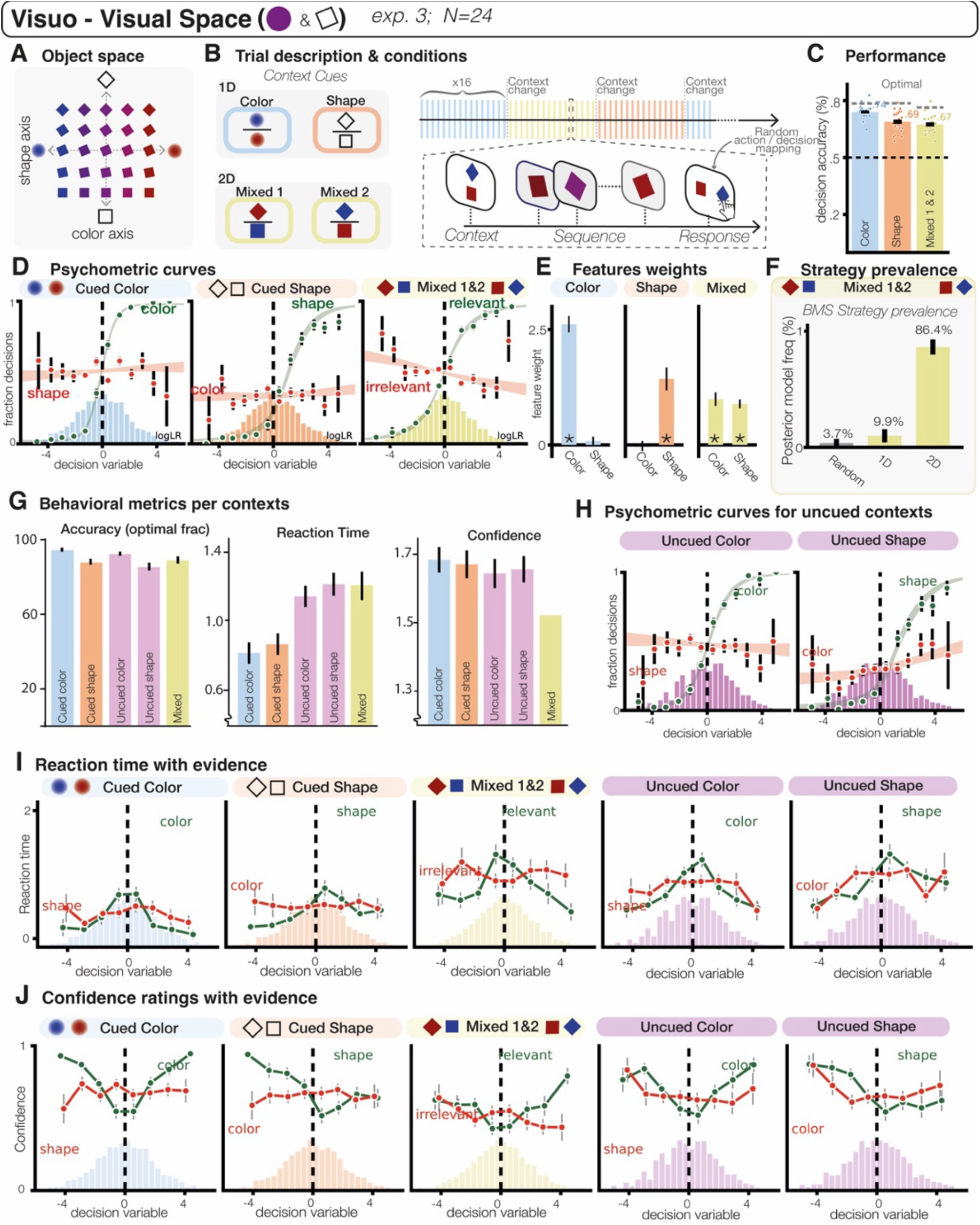
Behavioral metrics. **(A)** Objects spanned a two-dimensional space varying in color and in shape for exp 3 **(B)** Left: context cues. Right: block and trial description. **(C)** Human categorization accuracy for each condition: color, shape and mixed (2D) contexts. **(D)** Behavioral metrics. Left: color context. Center: shape context. Right: mixed (2D) contexts. Psychometric curves showing the relation between the decision variable (the accumulated category evidence in the active context) for the relevant (green) and irrelevant (red) feature, and the fraction of decisions. Dots and error bars indicate human data (mean ± s.e.m.) and shaded areas indicate psychometric fits (mean ± s.e.m.). **(E)** Left: color context. Center: shape context. Right: mixed (2D) contexts. Weight of each feature for the decision of the subjects in each condition. **(F)** Comparison of strategy prevalence in 2D conditions between a random strategy, a 1D strategy answering alternatively solely based on one or the other feature and a 2D strategy in which both features contribute to every decision. **(G)** Accuracy, Reaction Time and Confidence for each condition of exp. 3 **(H)** Psychometric curve for the two uncued conditions. **(I)** Similar analysis on reaction time revealed than participants were faster for larger amount of categorical evidence on the relevant axis, but RTs were independent from evidence on the irrelevant axis. **(J)** Similarly, participants confidence ratings were highest for larger amount of categorical evidence on the relevant axis, but independent from evidence on the irrelevant axis.

**Figure S2:**
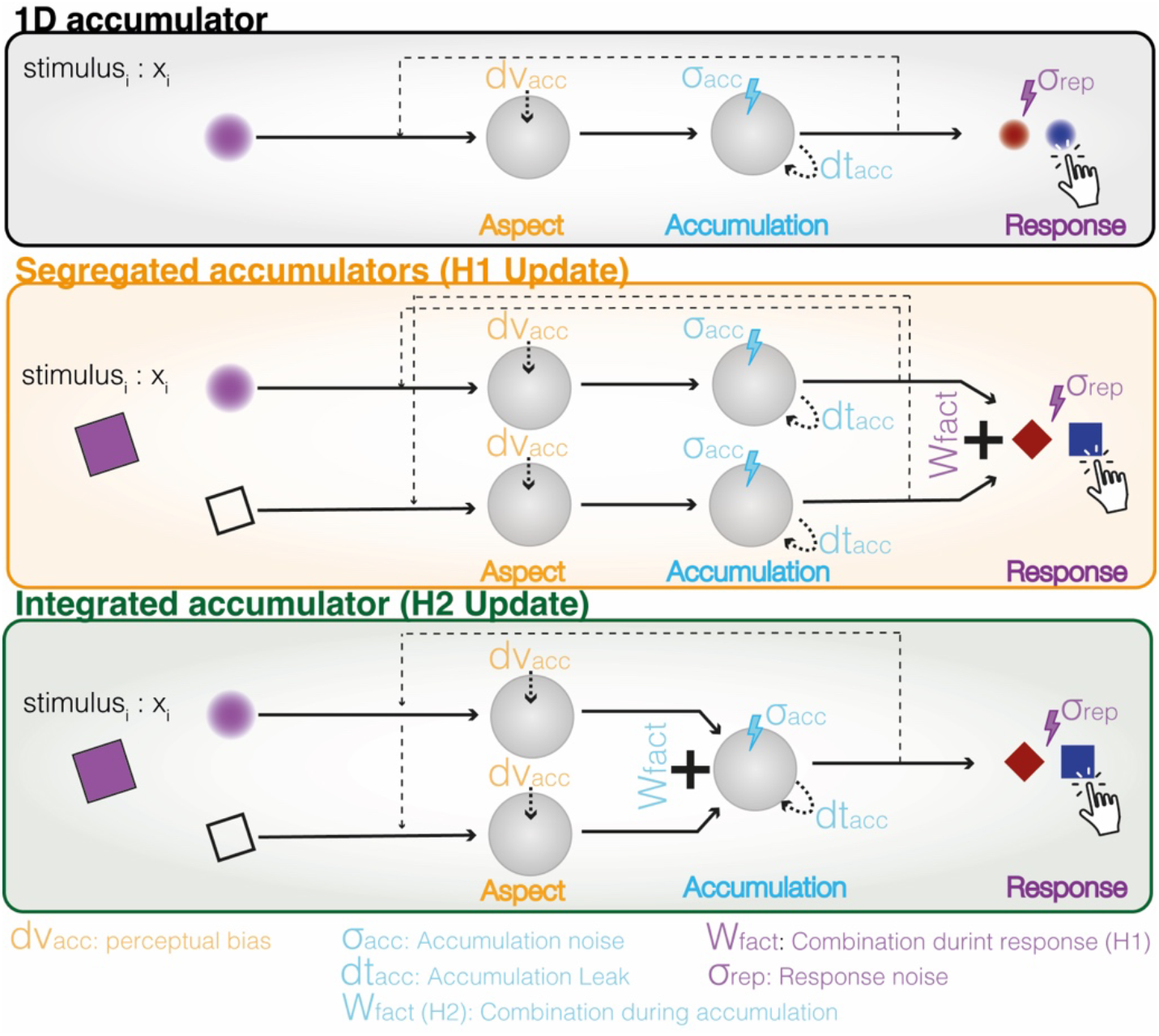
Cognitive computational model of behavior. *Top*: description of the accumulation model for a single (here color) feature. *Bottom:* Description of the two cognitive models of the two accumulation hypotheses. H1 postulates segregated accumulators for color and shape independently with the integration downstream of the accumulation, at the decision stage. On the other hand, H2 postulates a single accumulation process on a combination of the color and the shape features. Both models integrate suboptimalities including accumulation noise (σ_acc_), stimulus distortion (dv_acc_), temporal leak favoring recent observation over distant ones (dt_acc_), and response noise (σ_rep_).

**Figure S3:**
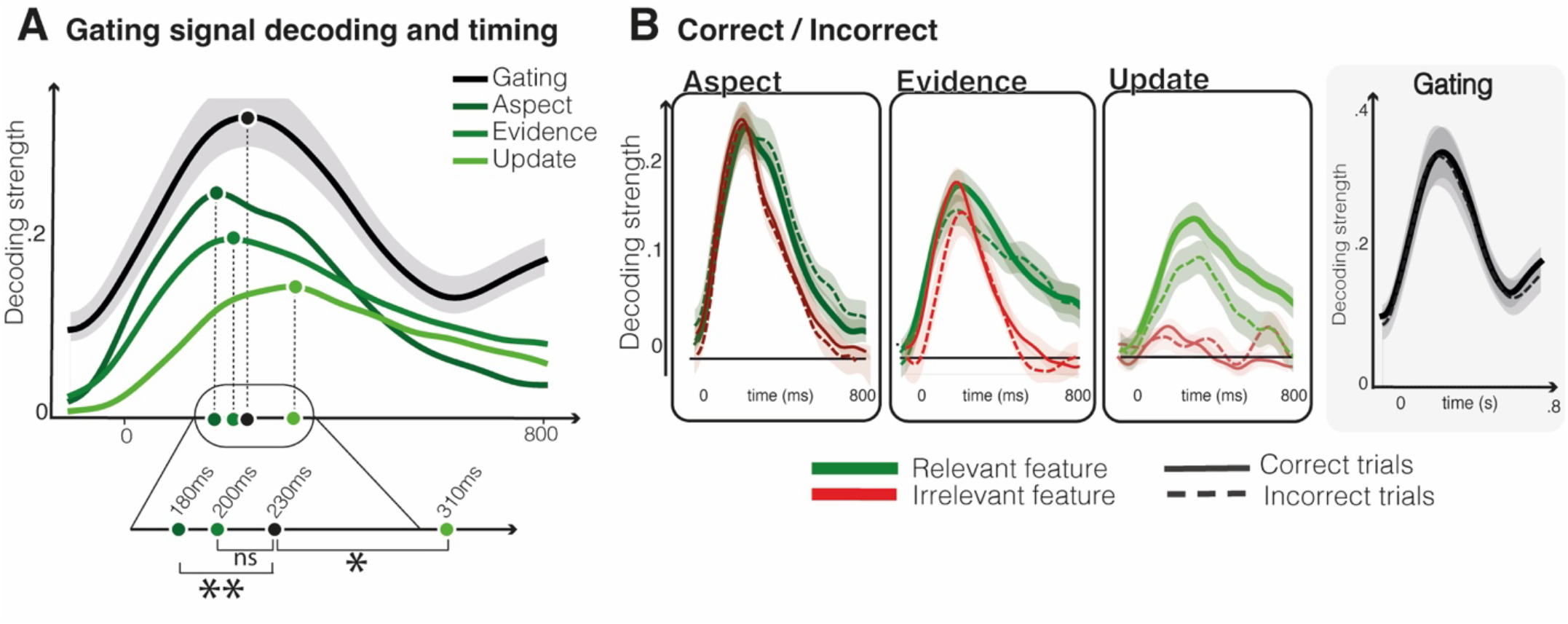
Timing of the different processing stage and gating of the irrelevant dimension. **(A)** We decoded the context (Color Vs Shape). While this context is constant over the course of the trial, we found that this code was reactivated after the presentation of each stimulus with a timing between its perception (aspect and salience) and its accumulation. This was interpreted as a gating signal, serving to project the multidimensional input into to select the relevant evidence only. To estimate significance between timing of the different stages of the processing hierarchy we performed bootstrap estimation of the maximum of each decoder (* p<0.05; **p<0.01; 1000 bootstrap resampling between subjects). **(B)**. Participants’ errors on such task could come from three distinct sources: 1. impaired perception, 2. impaired accumulation or 3. imperfect attentional gating. To disentangle between those three hypotheses, we spitted all decoders’ accuracies from correct and incorrect trials. The only difference observed was for the accumulation for the relevant feature (cluster p<0.01), favoring the second hypothesis over the first. Moreover, we found no modulation of the accumulation decoding accuracy for the irrelevant feature, suggesting that errors do not come from inoperative gating error and inopportune accumulation of the irrelevant dimension. Additionally, we found no modulation of the gating signal, speaking against context-dependent selection as a source of error.

**Figure S4:**
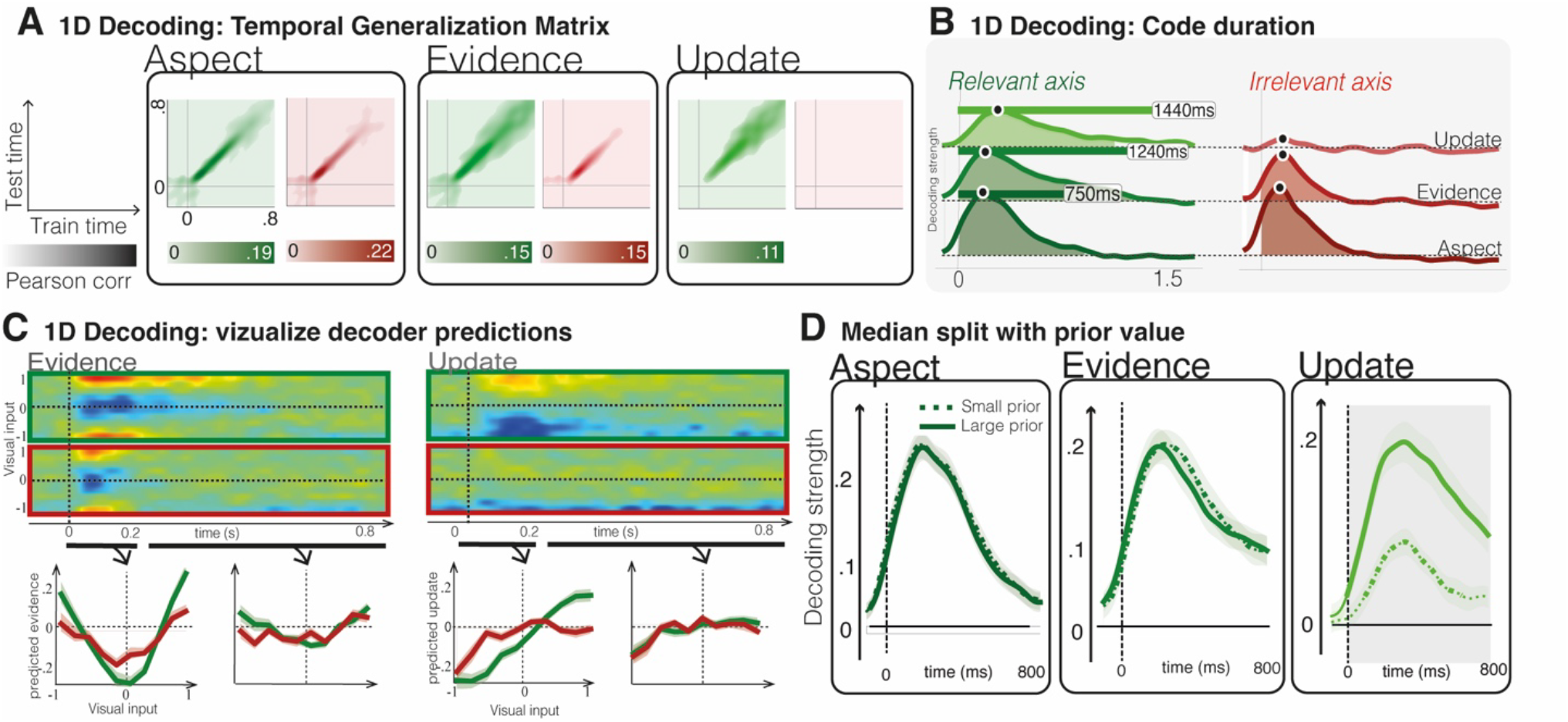
Complementary analyses of 1D decisions. **(A)** Generalization Across Time (GAT) matrices for the aspect, salience, and accumulation decoders, computed separately on the relevant and irrelevant axes for the Shape and Color contexts. **(B)** To assess the temporal persistence of each neural code, we repeated the decoding analyses for aspect, salience, and accumulation on the focused and ignored axes, extending the time window up to 1.6 s after stimulus onset. **(C)** For the salience and accumulation decoders, instead of quantifying performance using Pearson correlation with the true values, we directly plotted the decoder outputs as a function of the visual input. **(D)** Decoding accuracy for aspect, salience, and accumulation on the relevant axis was further divided using a median split based on prior belief (i.e., accumulated belief after element *i–1*). This analysis confirmed that lower prior belief (i.e., less information about the category before the current item) was associated with reduced accumulation update at the subsequent element.

**Figure S5:**
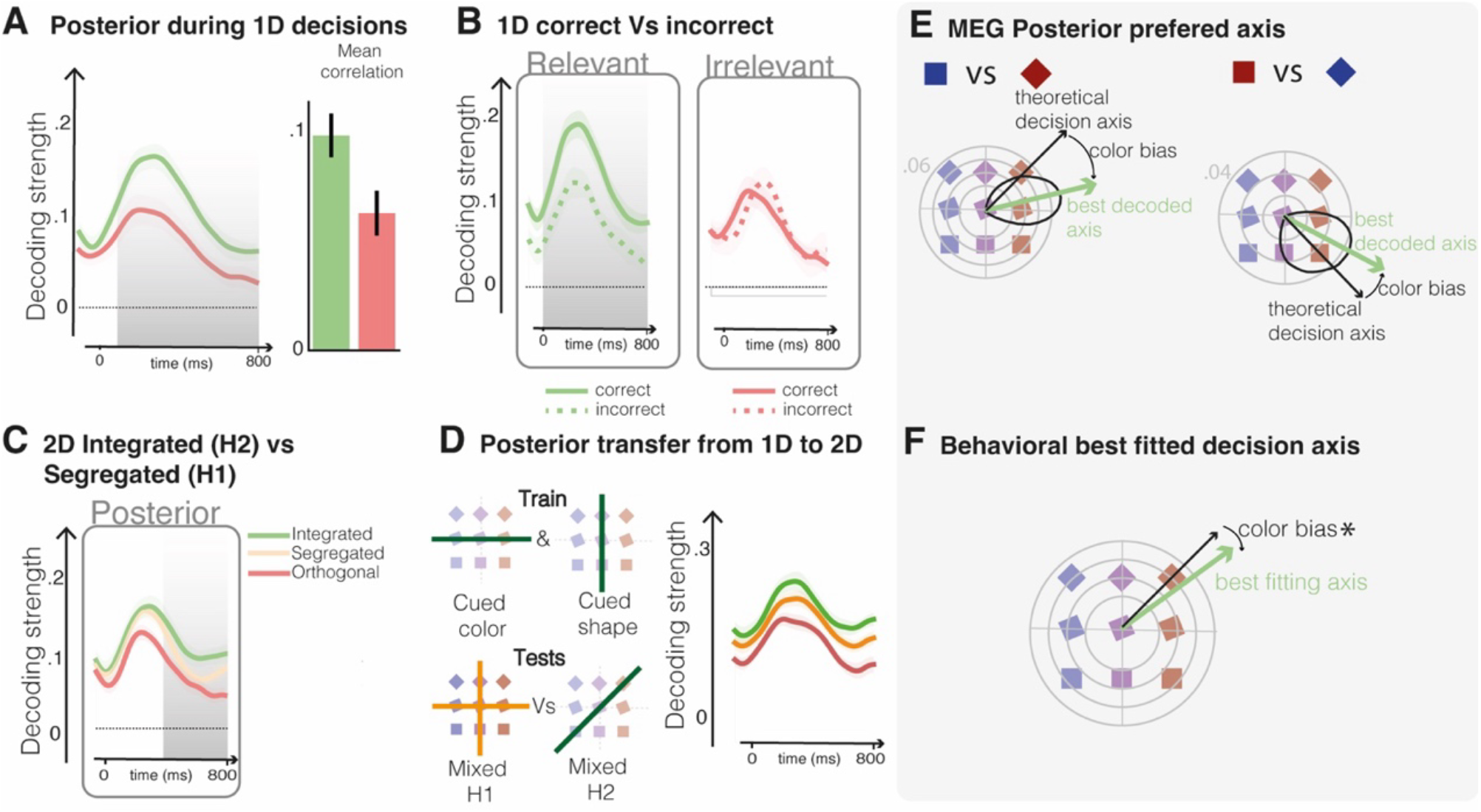
Neural decoding of posterior. In addition to the three primary metrics examined in this study (aspect evidence and update), we also included absolute value of the total evidence accumulated as a fourth metric (Posterior - P_i_). The posterior at time *i* corresponds to the total accumulated belief up to element *i* in the sequence. However, this additional metric presents two limitations. First, P_i_ is intrinsically correlated with P_i−1_. Second, because sequences are drawn from a normal distribution with an offset toward a given category, P_i_ systematically diverges as *i* increases (i.e., accumulated evidence grows with sequence length). Consequently, this measure cannot be interpreted as reflecting accumulated evidence alone, as it may also partially capture the position of the stimulus within the sequence. For this reason, we used this metric only as confirmatory support for previously demonstrated effects. **(A)** Decoding of posterior belief for the 1D contexts (shape and color) along the relevant and irrelevant axes revealed a strong selection effect for the relevant feature. **(B)** Only posterior decoding along the relevant feature was modulated by trial accuracy; decoding along the irrelevant feature was not. **(C)** In the 2D (mixed) contexts, we replicated the combination effect: posterior decoding accuracy was higher along the relevant axis than along the segregated feature axes or the orthogonal null axis. This finding further supports the hypothesis that humans preferentially accumulate combined feature evidence rather than processing the two features independently. **(D)** We also reproduced the transfer analysis: training the decoder on the segregated axis in the 1D contexts and testing it on the mixed contexts provided additional support for the combination hypothesis. **(E)** In the mixed contexts, we trained and tested the posterior decoder across all possible projection axes. The highest mean decoding accuracy was obtained for an axis combining the two features, but biased toward weighting color evidence more heavily than shape evidence. **(F)** This color bias is consistent with the stronger weights for color estimated in the mixed context (see also **Figure 1D**).

**Figure S6:**
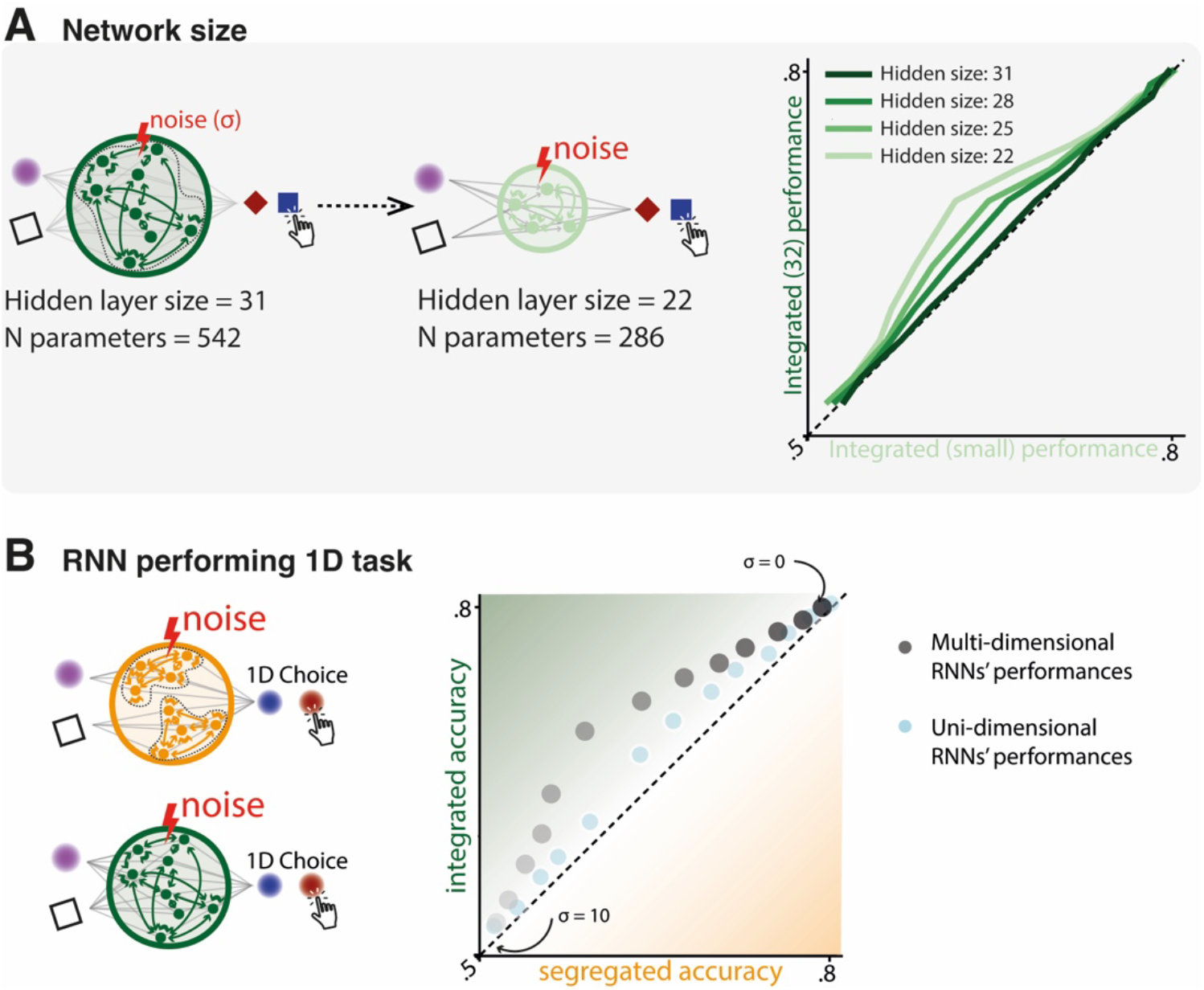
Complementary analyses of segregated vs integrated RNNS. **(A)** In the original analysis, sparse networks contained slightly more trainable parameters than split networks. Although the number of hidden-to-hidden parameters was identical due to matched overall sparsity **N**_**hh**_ = **size**_**hh**_^**2**^ / **2** the number of input-to-hidden parameters differed between architectures. Specifically, the split network had **N**_**ih**_ = **2** * **size**_**ih**_/**2** whereas the sparse network had **N**_**ih**_ = **2** * **size**_**ih**_. For RNNs with 32 hidden units, this resulted in a total of 544 trainable parameters for the split network compared to 576 for the sparse network. To determine whether this modest difference accounted for the greater noise robustness observed in split networks, we evaluated smaller sparse networks and compared them to the 32-unit non-split model. The sparse architecture with the closest parameter counts to the 32-unit split network was a 31-unit non-split network (542 parameters). We observed no difference in noise robustness between the 32-unit and 31-unit sparse networks, indicating that parameter count alone does not explain the robustness advantage of split networks. We further conducted a systematic analysis of smaller sparse networks (31, 28, 25, and 22 hidden units). Noise robustness decreased progressively with network size. Notably, the 32-unit split network (544 parameters) exhibited a level of robustness comparable to that of a 22-unit sparse network (286 parameters). These results rule out total parameter count as the primary determinant of the enhanced noise robustness observed in split networks. **(B)** We also replicated the noise robustness analysis using split and sparse RNNs trained on the 1D (Color) task. The sparse network was slightly more robust to noise. However, the magnitude of this effect was very small compared to the robustness differences observed in the 2D task. This contrast further supports the conclusion that the key factor underlying the robustness effect is the combined versus segregated accumulation of evidence in the 2D decision process, rather than architectural differences per se.

**Figure S7:**
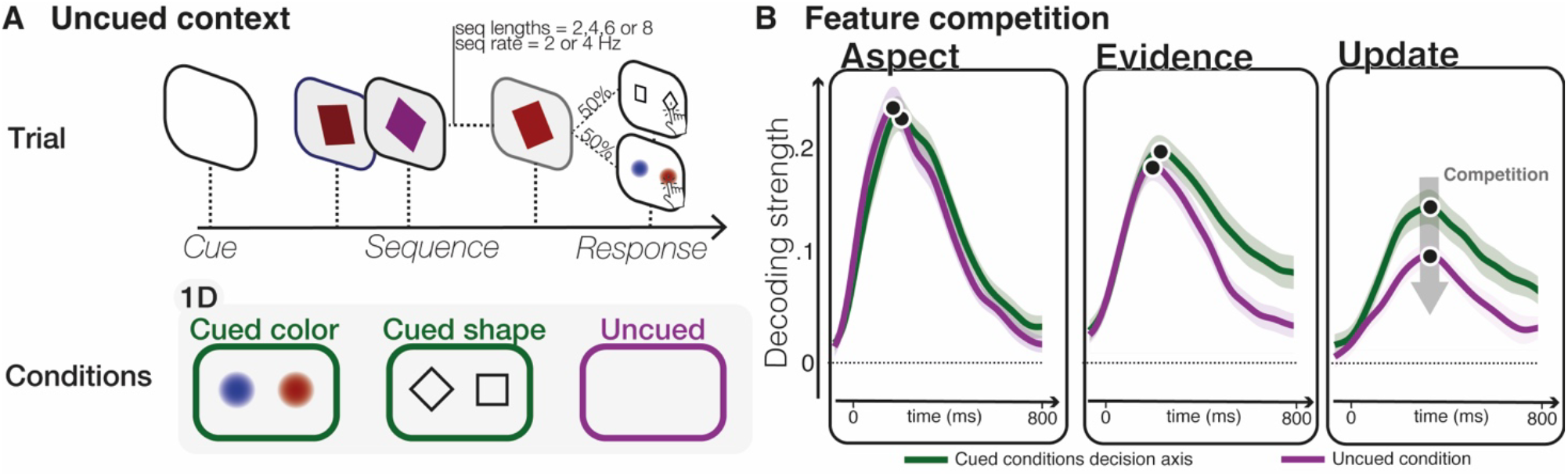
Competition effect during 1D accumulation. **(A)** During uncued trials, participants didn’t know in advance over which 1D category (color or shape) they would be prompted to make a decision. Accurate decision-making thus requires the accumulation of both color and shape in parallel. **(B)** Comparing the decoding accuracies between the cued and uncued contexts revealed that the accumulations of the two features were in competition, resulting in a lower encoding of each feature accumulation. This results further reinforce the idea that combination rather than segregated accumulation allows to mitigate the cognitive bottleneck imposed by the accumulation process, effectively avoiding the competition effect.

**Figure S8:**
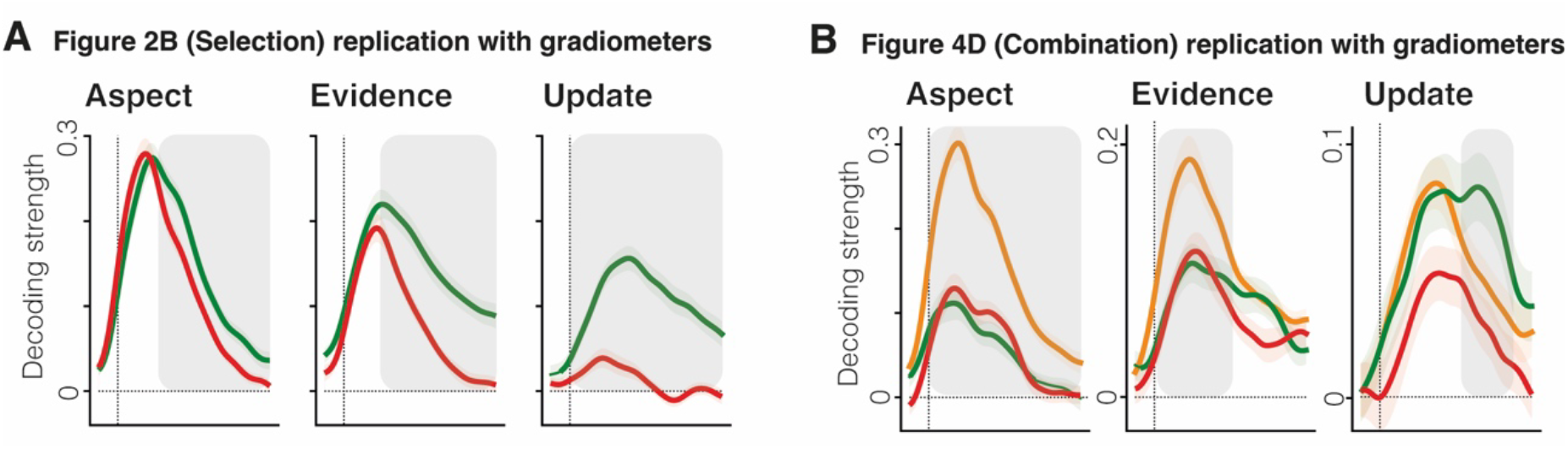
Replication of the MEG decoding analysis with gradiometers. All decoding analyses from the main text have been performed on the magnetometer sensors. To assess the robustness of our main effects, we recomputed all major analysis from **Figure 2B (A)** and **Figure 4D (B)** with gradiometers and replicated all findings.

**Figure S9:**
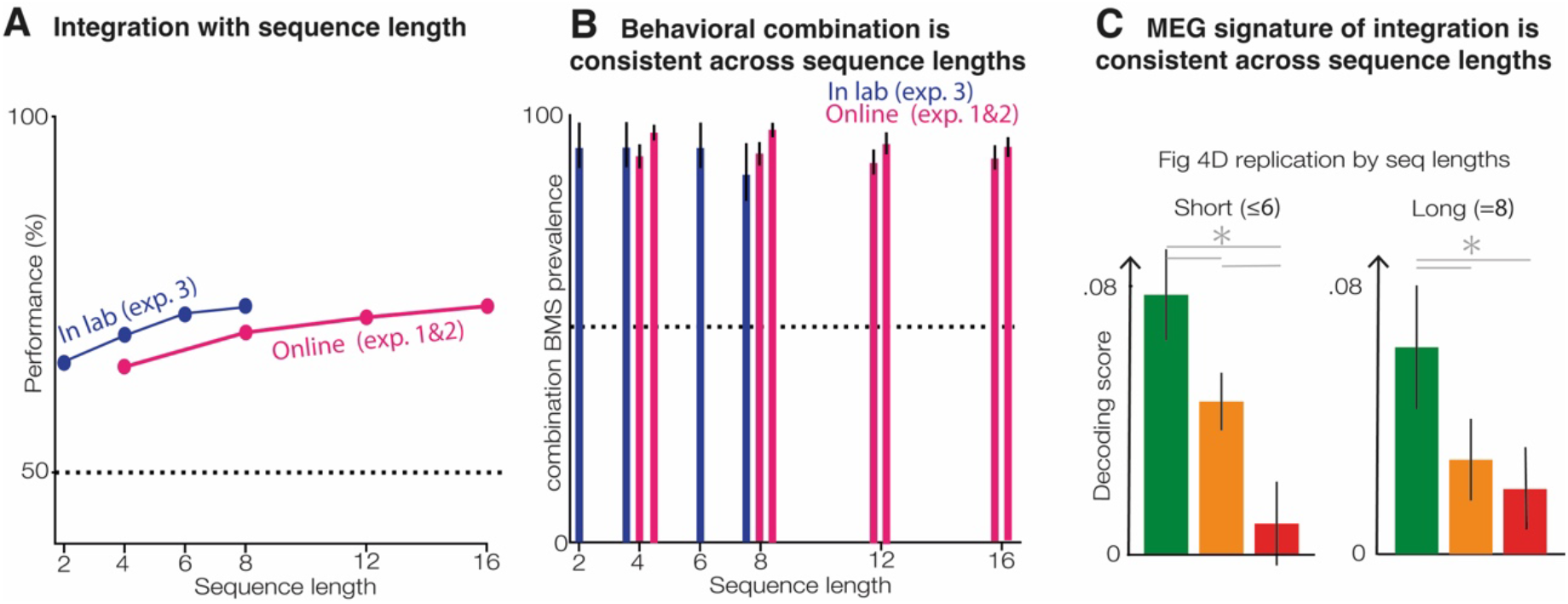
Exploring the effect of sequence lengths. **A**. Accuracy increases with sequence length, revealing efficient accumulation of evidence over multiple elements of the sequence. **B**. Strategy prevalence analyses (Fig1F/L & FigS1F) by sequence lengths reveals that the combination strategy prevalence is independent of sequence length? **C**. Replication of the integrated vs segregated analyses of the update in 2D decision (Fig 4D), for short (=<6 items) and long (=8 items) sequences. The pattern is independent of sequence length.

